# Novel mtDNA Imparts the Connective Tissue Disorder of a Tourette Pedigree

**DOI:** 10.1101/2022.02.25.481696

**Authors:** Patrick M. Schaefer, Leonardo Scherer Alves, Maria Lvova, Jessica Huang, Komal Rathi, Kevin Janssen, Arrienne Butic, Tal Yardeni, Ryan Morrow, Marie Lott, Kierstin Keller, Benjamin A. Garcia, Clair A. Francomano, Douglas C. Wallace

## Abstract

Mitochondrial dysfunction is associated with a range of clinical manifestations including neuropsychiatric and metabolic disorder. Here, we reanalyzed a family with an L-Histidine Decarboxylase (HDC) variant previously linked to Tourette syndrome but with associated connective tissue and metabolic features of unknown etiology. We identified a mitochondrial haplogroup J-defining mutation on the haplogroup H background that functionally interacts with the L-Histidine Decarboxylase variant via calcium homeostasis. Our findings establish how a common mtDNA variant on a different mtDNA background can result in mitochondrial dysfunction, demonstrate a role for histaminergic signaling in modifying mitochondrial phenotypes, and link mitochondria dysfunction to connective tissue phenotypes.

---

Mitochondria are increasingly recognized as integral factors in many pathologies, including diabetes or neuropsychiatric disorders (1). While most components of the mitochondria are nuclear-encoded, the maternally inherited mitochondrial DNA (mtDNA) still retains 13 essential subunits of oxidative phosphorylation (OxPhos) as well as the structural RNAs for the mitochondrial protein synthesis machinery (2–4). During human evolution, functional mtDNA variants arose along radiating maternal lineages that resulted in subtle changes in mitochondrial bioenergetics. These permitted humans to adapt to the new environments they encountered as they migrated out-of-Africa and around the globe. These adaptive variants function within the context of pre-existing mtDNA functional variants resulting in an integrated bioenergetic state. Those variant combinations that proved regionally beneficial became enriched to form geographically constrained groups of related haplotypes, known as a haplogroup (5).

Tourette syndrome is an autism spectrum disorder (ASD) characterized by motor and vocal tics with a strong genetic component, and a mtDNA mutant has been shown to produce ASD-like symptoms in a mouse model (6). In a large two-generation pedigree, linkage analysis identified a null variant in the L-Histidine Decarboxylase (HDC) gene (W317X) as a candidate for causing Tourette syndrome (7). However, this pedigree manifests several unusual characteristics that cannot be explained by Mendelian inheritance of the HDC gene variant. First, a paternal heterozygous variant was passed to all 8 living offspring, while seven other conceptions were lost as miscarriages. The probability of this chain of events to happen by chance is 0.4%. Second, the mother and all living offspring display a complex metabolic phenotype in association with a skeletal and connective tissue disorder that resembles Ehlers-Danlos syndrome.

We have tested the hypothesis that this pedigree involves a deleterious maternally-inherited mtDNA for which the HDC variant was protective. This resulted in only the offspring that inherited the HDC variant surviving, while still manifesting the metabolic and connective tissue defects associated with the deleterious mtDNA.

## Results

### Family harbors a dysfunctional mtDNA variant

We reevaluated, assembled, and summarized the clinical data for the two-generation pedigree of Tourette syndrome and obtained additional data from subsequent family assessments (7). In addition to a range of neurological symptoms (Tourette, Obsessive-Compulsive Disorder (OCD), ASD) reported in the original paper (7), additional information was identified for the mother (Fig. 1a, I2) and her offspring. This elaborated on the previously reported information pertaining to skeletal and connective tissue manifestations, including Chiari malformation, cranial-cervical instability, hypermobility of joints, skin herniation, wound dehiscence, aortic valve disease, ankylosing spondylitis, diabetes, fatigability, and dysautonomic manifestations (see Appendix). The apparent matrilineal transmission of this complex array of clinical manifestations in all 8 offspring suggested a mtDNA disorder. Our resulting interpretation of the two-generation pedigree is summarized in Fig. 1a.

**Fig. 1.**
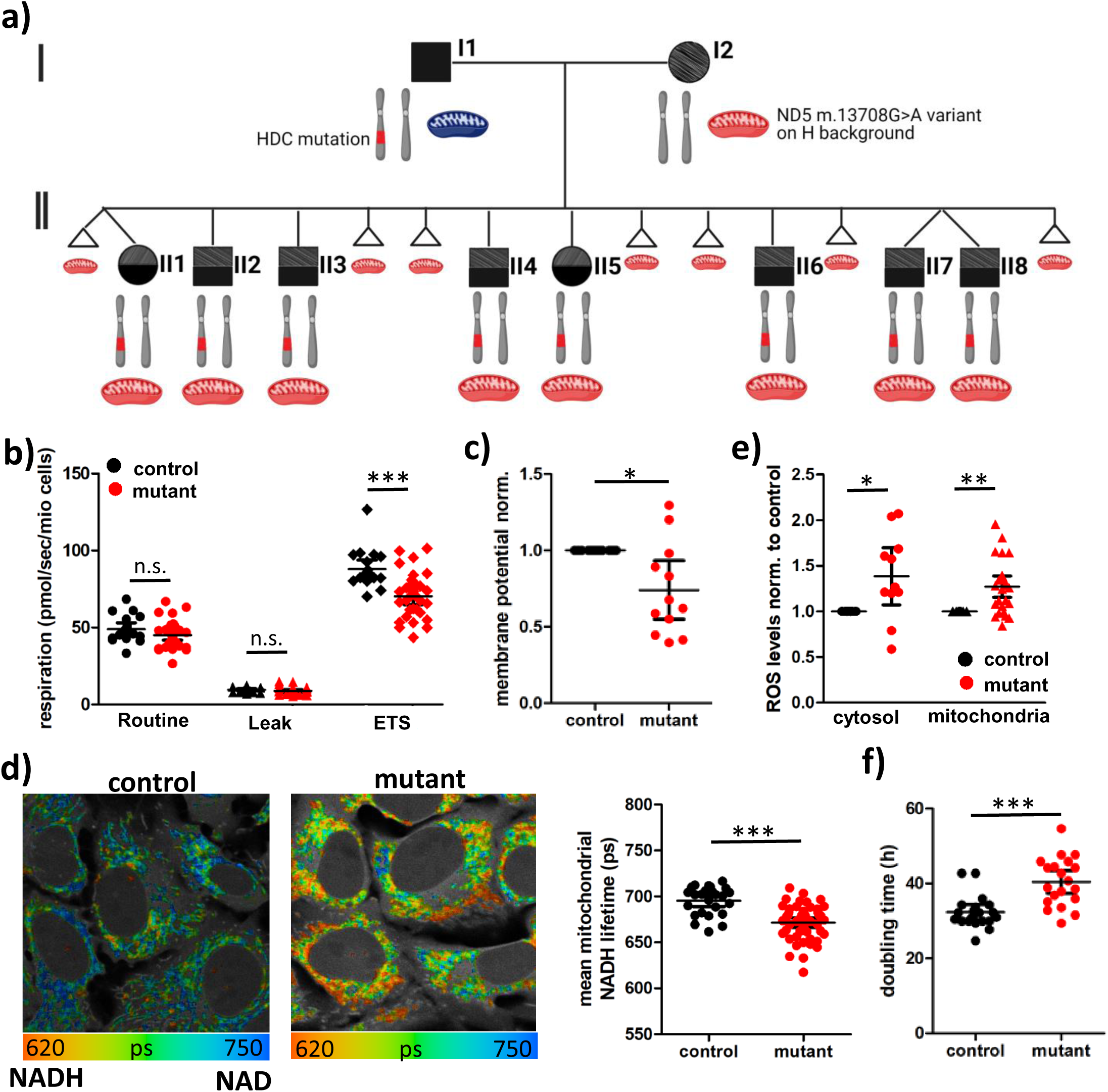
*ND5* m.13708G>A-H7 mtDNA reduces bioenergetic function. **a)** Two generation pedigree with squares indicating males, circles females and triangles miscarriages. Black color indicates Tourette syndrome phenotype, grey-shaded indicates connective tissue phenotype. Chromosomes indicate the genotype for chromosome 15, with the father and all children being heterozygous for the W317X HDC variant. Red mitochondria indicate maternal mitochondria harboring the *ND5* m.13708G>A variant on haplogroup H7 background. **b)** Respirometry of intact transmitochondrial cybrids harboring control (black) or patient (mutant, red) mitochondria (control: n = 19, mutant n = 31, technical duplicates, Mann Whitney test). **c)** Mitochondrial membrane potential in control and mutant cybrids measured as red/green fluorescence of JC-1 quantified by flow cytometry normalized to control (n=12, technical duplicates, paired t-test). **d)** Fluorescence lifetime imaging microscopy (FLIM) of mitochondrial NADH in control and mutant cybrids. False-color coding of NADH fluorescent lifetime with blue indicating a longer lifetime (more oxidized NAD^+^/NADH) and red indicating a shorter lifetime (more reduced NAD^+^/NADH). Grey indicates non-mitochondrial autofluorescence. Quantification of the mean mitochondrial NADH lifetime (n = 5 with 5 image sections and >25 cells/independent experiment, unpaired t-test). **e)** Cytosolic and mitochondrial ROS levels in control and mutant cybrids quantified as the fluorescence intensity of DCDFA (cytosol, n = 11 in technical duplicates, paired t-test) or Mitosox (mitochondrial, n= 26 in technical duplicates, paired t-test) normalized to control. **f)** Doubling time of control and mutant cybrids (n=20, Wilcoxon signed rank test). Error bars display 95% confidence intervals (CI). Significances are indicated by stars with * = p<0.05, ** = p<0.01, *** = p<0.001.

To determine if there was a mtDNA contribution to this pedigree, we sequenced the mtDNA of the mother and five of her children, their mtDNA variants being presented in Fig.S1a. The mtDNA haplogroup is H7, but this family’s mtDNA also contains the haplogroup J-defining variant *ND5* m.13708G>A (codon 458 Alanine to Threonine). This is noteworthy since the *ND5* m.13708G>A is very rarely associated with haplogroup H7 (AF = 0.86%, 3/348) in Mitomap (8) and rarely on other haplogroup H mtDNAs (Fig.S1b, c). To assess whether the *ND5* m.13708G>A variant on haplogroup H7 affects mitochondrial function, we created transmitochondrial cybrids that have the same 143B(TK^-^) cell nDNA but harbor either the patient mtDNA (mutant) or the mtDNA of a haplogroup-matched normal control (control) (Fig.S1a, Fig.S2a). We observed a lower electron transport system capacity with reduced complex I and II respiration in the mutant cybrids (Fig. 1b, Fig.S2b, d). In line with the lower respiration, the mutant cybrids displayed a reduced mitochondrial membrane potential (Fig. 1c) and NAD^+^/NADH redox ratio (Fig. 1d). In addition, both mitochondrial and cytosolic ROS levels were increased (Fig. 1e, Fig.S2e), doubling time was longer (Fig. 1f) and apoptosis was increased (Fig.S2f) in the mutant cybrids. In contrast, haplogroup J cybrids harboring the *ND5* m.13708G>A variant in its standard context showed similar respiration haplogroup H cybrids and even reduced ROS production (Fig.S3). Hence, when arising in the wrong context, the *ND5* m.13708G>A variant is incompatible with the resident haplogroup H mtDNA functional variants resulting in mitochondrial dysfunction and clinical manifestations.

To further evaluate the impact of the *ND5* m.13708G>A-H7 mtDNA mutant on cellular physiology, we performed multi-omics on the cybrids. This revealed profound changes to the gene expression, histone acetylation, and metabolite profile (Fig. 2).

**Fig. 2.**
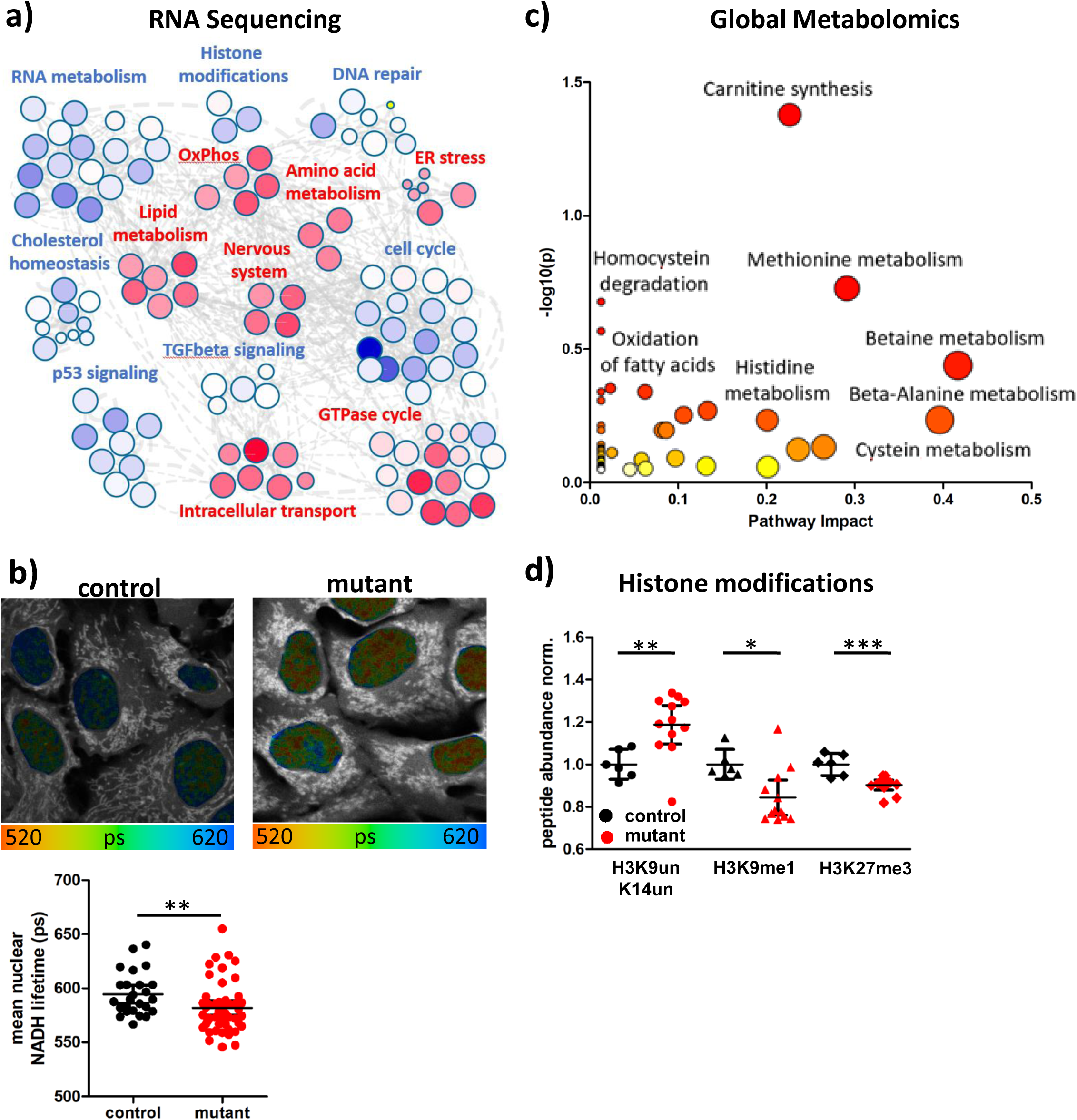
*ND5* m.13708G>A-H7 mtDNA profoundly alters metabolite and expression profile. **a)** Cluster analysis from GSEA of RNASeq in mutant versus control cybrids using Cytoscape. Every node represents a significant pathway with blue indicating upregulation and red downregulation of the pathway in variant versus control. **b)** Fluorescence lifetime imaging microscopy (FLIM) of nuclear NADH in control and mutant cybrids. False-color coding of NADH lifetime with blue indicating a longer NADH fluorescent lifetime and a more oxidized NAD+/NADH redox ratio and red indicating a shorter NADH fluorescent lifetime or more reduced redox ratio. Grey indicates non-nuclear autofluorescence. Quantification of the mean nuclear NADH lifetime revealed a more reduced redox ratio in mutant cybrids (n = 5 with 5 image sections and >25 nuclei/independent experiment, unpaired t-test). **c)** Metabolic pathway analysis of global untargeted metabolomics using significantly (p < 0.05) altered metabolites between mutant and control cybrids. Size of the dots indicates pathway impact (x-axis) and color (white – yellow – red) indicates p-value with red corresponding to higher significance. **d)** Selected histone modifications of histone 3 in control and mutant cybrids displayed as peptide abundance normalized to control. (n = 6, 12, unpaired t-test).

RNA sequencing revealed downregulation of OxPhos and nicotinamide metabolism in mutant cybrids (Fig. 2a, Fig.S4). This goes along with a more reduced nuclear NAD^+^/NADH redox ratio (Fig. 2b) and altered nucleotide levels and TCA cycle intermediates in the mutant cybrids (Fig.S4b, c). Interestingly, while nicotinamide levels were increased, 6-methylnicotinamide was significantly decreased in the mutant cybrids (Fig.S4b), indicating a problem in the SAM-dependent degradation of nicotinamide. Global metabolomics point to alterations in methionine metabolism and homocysteine degradation (Fig. 2c) and RNA sequencing revealed a downregulation of the one carbon metabolism (Fig. 2a, Fig.S4d). Targeted metabolomics verified reduced homocysteine levels but increased cystathionine levels and trends towards a reduced SAM/SAH and GSSG/GSH ratio in the mutant cybrids (Fig.S4e), revealing a defect in the mitochondrial one carbon metabolism. Consistent with lower methyl-donor availability, we detected histone hypomethylation in the mutant cybrids (Fig. 2d, Fig.S5). In addition, global metabolomics revealed increased levels of acylcarnitines (Fig. 2c, S6d) in the mutant cybrids, indicating alterations in fatty acid metabolism. This is further supported by significant changes to the ketone body metabolism and an upregulation of the cholesterol biosynthesis pathway in RNA sequencing (Fig.S6a,b). Importantly, RNAseq revealed significant differential expression of genes associated with collagen formation and neurodevelopment in the patient mtDNA cybrids (Fig. 2a, Fig.S6c), linking the mitochondrial variant with the connective tissue phenotype and Tourette syndrome of the family.

Taken together, the *ND5* m.13708G>A variant on the haplogroup H7 background affects mitochondrial fatty acid metabolism, one carbon metabolism, nicotinamide metabolism and collagen metabolism, thus profoundly changing cellular gene expression and physiology.

### HDC variant acts as a beneficial nuclear modifier via calcium homeostasis

Next, we asked whether the heterozygous HDC variant from the father could have a beneficial effect on the maternal *ND5* m.13708G>A-H7 mtDNA, thus explaining why all living offspring carry the HDC variant while seven others died *in utero*. Given the central role of HDC in histamine signaling (9) and subsequent calcium release from the ER (10), we assessed calcium homeostasis in the cybrids.

We found higher cytosolic calcium but reduced mitochondrial calcium levels in the mtDNA mutant cybrids at baseline (Fig. 3a). Histamine-induced calcium release from the ER resulted in a stronger increase of cytosolic calcium in the mutant cybrids (Fig. 3b), which was weakened but retained upon pretreatment with a mitochondrial calcium uptake inhibitor (Fig.S7c) but not after thapsigargin-induced calcium release (Fig.S7d). This demonstrates an increased histamine sensitivity in the mutant cybrids, which corresponds to an upregulation of the histamine 1 receptor signaling pathway on the RNA level (Fig.S7e). At the same time, the partial rescue with the MCU inhibitor indicates a lower mitochondrial calcium uptake in the mutant cybrids. This is confirmed by a lower increase of mitochondrial calcium upon histamine treatment (Fig.S7b) and downregulation of the mitochondrial calcium ion transport (Fig.S7a) in the mutant cybrids. Taken together, the *ND5* m.13708G>A variant on haplogroup H7 background alters calcium homeostasis and histamine signaling, providing a link to the HDC variant.

**Fig. 3.**
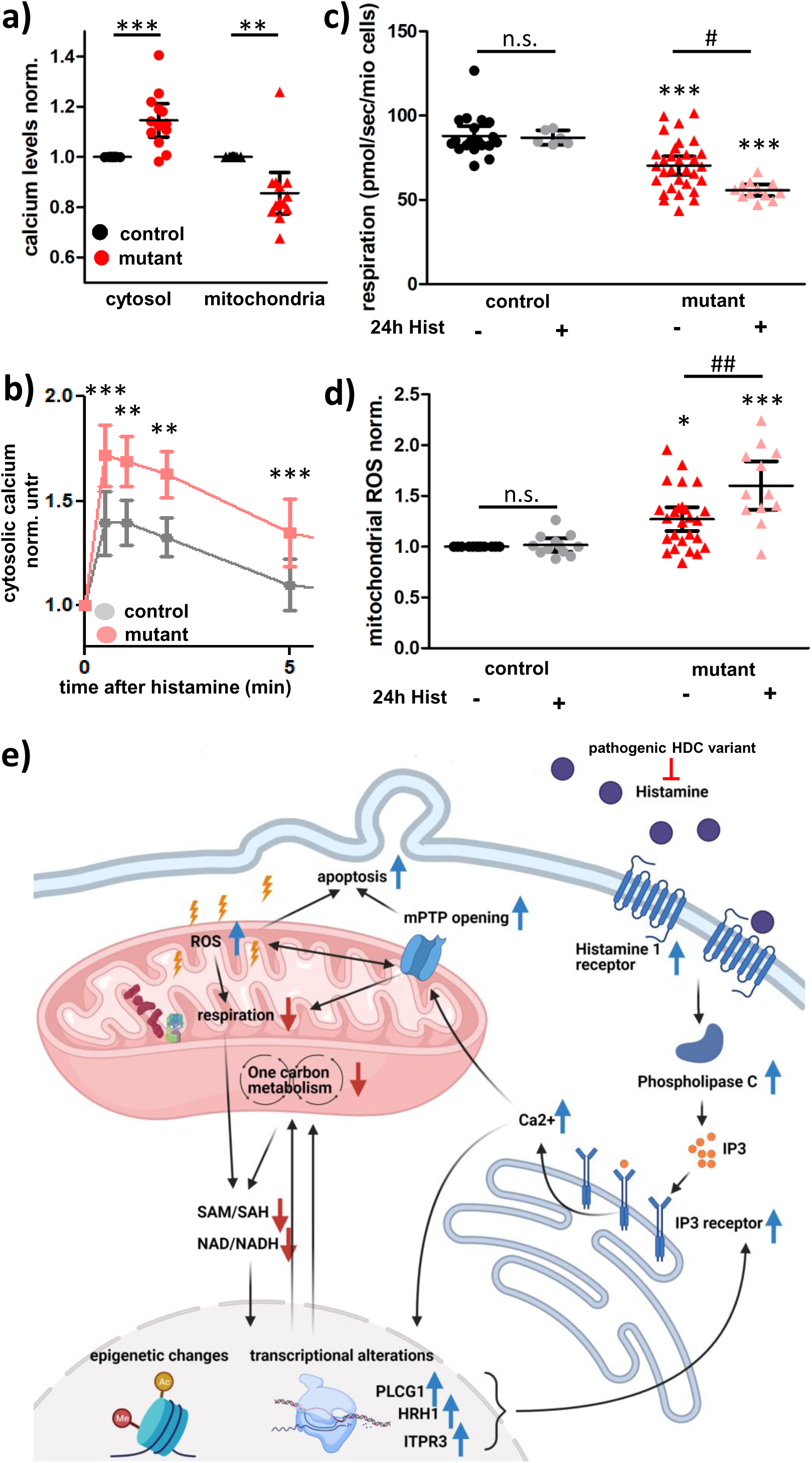
Calcium homeostasis links paternal HDC variant with the *ND5* m.13708G>A-H7 mtDNA. **a)** Cytosolic and mitochondrial calcium levels in control and mutant cybrids normalized to control measured by confocal microscopy or flow cytometry using Fura Red (cytosol, n = 6 microscopy, n = 7 flow cytometry, paired t-test) or Rhod-2 (mitochondria, n = 13 flow cytometry, paired t-test). **b)** Cytosolic calcium levels in mutant and control cybrids in response to a 100 µM histamine stimulus, normalized to calcium levels pre-addition (time point 0). Measured using Fura Red and confocal microscopy (n = 7, paired t-test). **c/d)** Electron transport system capacity (c) and mitochondrial ROS levels (d) of control and mutant cybrids with and without pretreatment with 100 µM histamine for 24h. Significances between mutant and control (*) and between treatments (#) were calculated using Kruskal Wallis test (n=19, 6, 31, 12, technical duplicates) for respirometry and One-Way ANOVA (n = 26, 12, 26, 12, technical duplicates) for ROS. **e)** Diagram visualizing the interaction between a pathogenic variant in the Histidine Decarboxylase and the *ND5* m.13708G>A-H7 mtDNA. Blue arrows indicate an increase and red arrows a decrease in the mutant cybrids. The *ND5* m.13708G>A-H7 mtDNA reduces respiration and one carbon metabolism, resulting in epigenetic and transcriptional changes that include an upregulation of Histamine 1 receptor (HRH1), Phospholipase C (PLCG1), and IP3 receptor (ITPR3). Upon histamine, this leads to increased calcium release from the ER, mitochondrial calcium overload, mitochondrial permeability transition pore (mPTP) opening, and apoptosis. The HDC variant reduces the formation of histamine from histidine, thereby preventing histamine toxicity. Created with BioRender.com.

To simulate an increased HDC activity, we exposed the cybrids to histamine for 24h. The m.13798G>A-H7 mutant but not control cybrids showed a further reduced respiration (Fig. 3c, Fig.S8a) and increased ROS production (Fig. 3d) after treatment with 100 µM histamine for 24 h. This could be blocked by simultaneous treatment with the histamine 1 receptor antagonist pyrilamine (Fig.S8c-e). In addition, we found an increased mitochondrial permeability transition pore (mPTP) opening in the mutant cybrids (Fig.S8b), indicating that a mitochondrial calcium overload induced via histamine 1 receptor-mediated calcium release from the ER contributes to mutant cybrid cell death (Fig. 3e). Pyrilamine inhibition of the histamine 1 receptor slightly reduced mitochondrial respiration in controls but mitigated the mitochondrial phenotypic difference between mutant and control cybrids (Fig.S9b-h). In summary, lower histamine levels due to the paternal HDC variant mitigate the increased calcium overload toxicity for the *ND5* m.13708G>A variant on the H7 background, thus explaining why all of the surviving offspring inherited the HDC mutant allele.

The remaining question is why the mother, who does not carry the HDC variant, was affected with the metabolic and connective tissue disorders but was still alive. One possibility is that the mother harbored a nuclear genotype that was protective of the *ND5* m.13708G>A-H7 variant mitochondrial defect. To assess if the mother’s genotype affected her mitochondrial physiology, we created immortalized lymphoblasts from the mother, 4 of her children and 3 haplogroup-matched controls. For the mother and her children, we found no changes in respiratory capacity or calcium handling compared to controls, though a switch from complex I to complex II respiration and increased cytosolic ROS was observed (Fig. 4a-d). This mild mitochondrial phenotype compared to the mutant cybrids might be explained by low and thus undetectable expression of HDC and histamine 1 receptor in the lymphoblasts, similar to histamine 1 receptor inhibition in cybrids (Fig.S9).

**Fig. 4.**
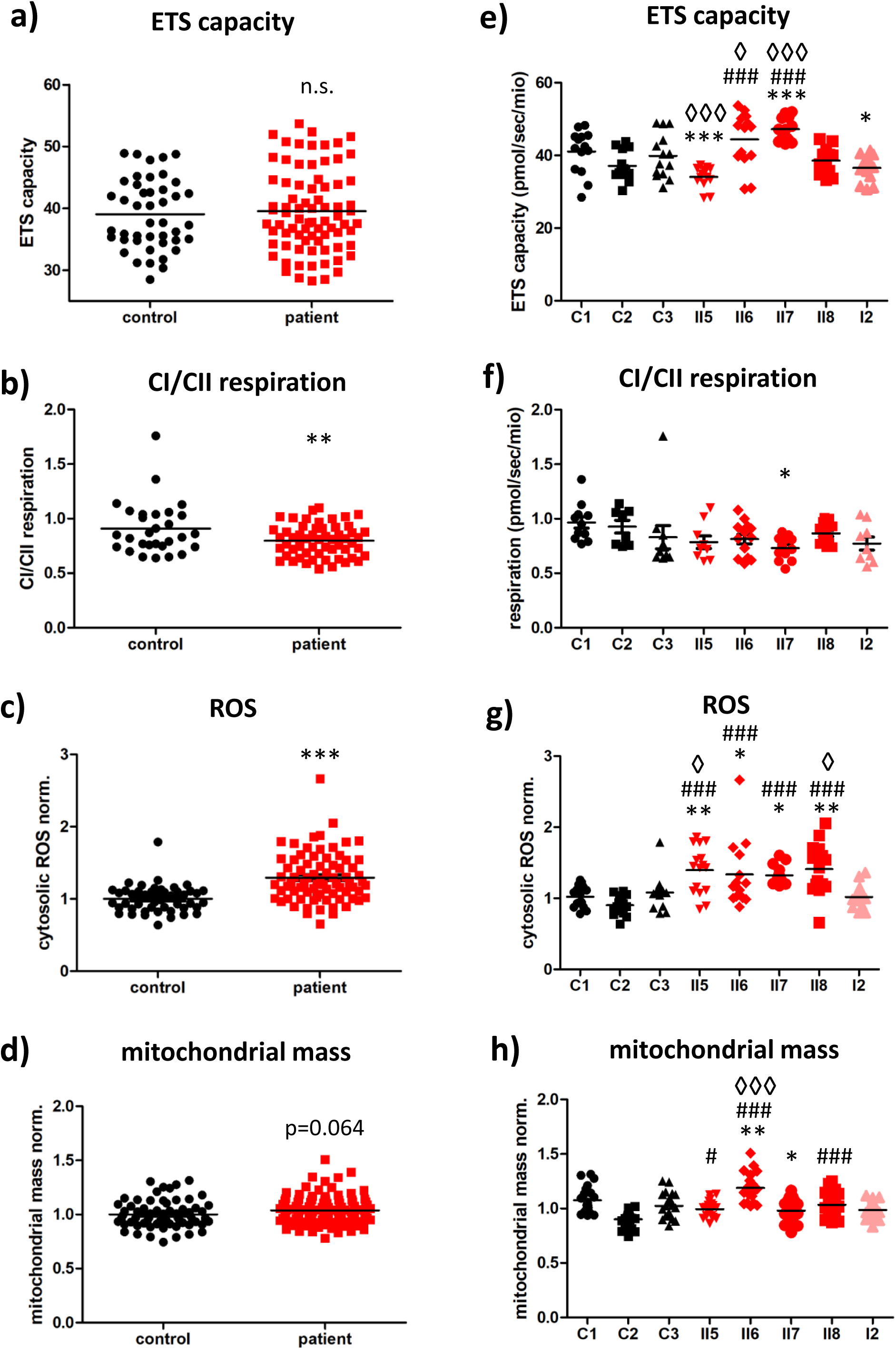
Patient lymphoblasts show increased ROS levels and a switch from complex I to complex II respiration. Patient lymphoblastoid cells from the mother (I2) and 4 offspring (II5-II8) are compared to 3 haplogroup-matched controls. **a/e)** Electron transport system capacity in intact control and patient lymphoblastoid cells (n = 14). **b/f)** Complex I/complex II respiration in permeabilized lymphoblastoid cells (n= 11,8,10,13,13,9,12,9) **c/g)** Cytosolic ROS levels in control and patient lymphoblastoid cells quantified as the fluorescence intensity of DCDFA normalized to control (n = 16,16,15,15,14,16,15,16). **d/h)** Mitochondrial mass of control and patient lymphoblastoid cells quantified as fluorescence intensity of Mitotracker CMX ROS or Mitotracker Deep Red in flow cytometry normalized to control (n = 11). Significance for patient versus control (a-d) was calculated using unpaired t-test and significances for individual patients versus individual controls (e-h) were calculated using One-way ANOVA or Kruskal Wallis test. Significances relative to control are indicated by * for C1, # for C2, and ◊ for C3.

Comparing the mother’s mitochondrial physiology to that of her children (Fig. 4e-h), we found that the lymphoblasts of all children but not the mother had increased ROS levels compared to control lymphoblasts (Fig. 4g, I2). Since increased ROS predisposes to apoptosis, the mother may harbor a multiple allele antioxidant genotype not transmitted to her offspring.

To verify if lowering ROS could mitigate the mother’s phenotype, we treated the *ND5* m.13708G>A-H7 cybrids with N-Acetyl-cysteine (NAC) and nicotinamide riboside (NR). Both partially rescued the mitochondrial defect in the mutant cybrids, resulting in a shorter doubling time, higher respiration, and lower cytosolic ROS levels (Fig.S10). This not only supports the mother harboring an enhanced antioxidant genotype but also suggests new therapeutic approaches for this family and perhaps other patients with mitochondrially-based connective tissue disorders.

## Discussion

We reanalyzed an unusual family with familial Tourette syndrome, metabolic dysfunction, and connective tissue disorder. Mendelian inheritance of a heterozygous HDC variant alone could not explain why all eight living children carry the HDC variant, suggesting that the seven additional miscarriages were the conceptuses that inherited the father’s normal HDC allele. In identifying an out-of-context mtDNA mutant in the patient family, we gained three crucial insights into the diversity of mitochondrial disorders and the mitochondrial etiology of the Tourette pedigrees patient phenotypes.

First, we could establish that a common haplogroup J-defining mtDNA variant causes mitochondrial dysfunction when combined with haplogroup H7, while the same variant in haplogroup J does not affect bioenergetics. This points to an incompatibility of *ND5* m.13708G>A variant with mtDNA variants present in haplogroup H7. Incompatibility between haplogroup-associated mtDNA variants may explain why the mtDNA is strictly maternally inherited, thus blocking inter-haplogroup recombination and the mixing of incompatible variant alleles (2). The existence of functional alterations in mtDNAs has been documented when mtDNAs of different origins have been mixed within a mouse and when different mtDNAs are placed on the same nDNA background (11–14). While the exact molecular basis of the intra-mtDNA *ND5* m.13708G>A haplogroup H7 incompatibility is unknown, one might hypothesize that mitochondrial electron transfer is hampered, resulting in lower respiration and higher ROS production. Taken together, we demonstrate that cross mtDNA mixing between common mtDNA variants can result in incompatibility and mitochondrial disorders. Clinically, this implies the need for not only checking the frequency of mtDNA variants but also the frequency of co-occurrence with other mtDNA variants when evaluating mtDNAs for pathological relevance.

Second, we demonstrate a mitochondrial-nuclear interaction with the paternal HDC variant and maternal mtDNA by establishing an important role for histaminergic signaling in modifying the mitochondrial phenotype. We showed that the HDC null allele (W317X) acts as a beneficial nuclear modifier for the deleterious maternal mtDNA. Given the crucial role for calcium signaling in embryogenesis (15), it seems plausible that the seven miscarriages were a result of a non-compensated dysregulation in calcium homeostasis due to the presence of the maternal mtDNA but absence of the HDC variant. While the mother herself does not harbor the HDC variant, we found lower cytosolic ROS levels in her lymphoblastoid cells compared to those of her children, suggesting the presence of additional nuclear antioxidant modifiers. Consistent with previous reports for mitochondrial dysfunction in cells (16) or mice (6) lowering the ROS levels in the *ND5* m.13708G>A-H7 cybrids rescued the mitochondrial phenotype.

Third, we revealed a putative mitochondrial etiology for the connective tissue phenotype of the patients and potential involvement in the Tourette syndrome. The connective tissue phenotype of the family resembles Ehlers Danlos Syndrome, which is commonly caused by mutations in collagen, other elements of the connective tissue, or the enzymes involved in processing the structural proteins of the extracellular matrix (17). It has been shown that collagen VI deficiency results in mitochondrial dysfunction, including calcium deregulation and mPTP opening in mice with a form of muscular dystrophy (18). Interestingly, mPTP desensitization improved the muscular dystrophy, suggesting a bidirectional relationship (19). Here we provide direct evidence that mitochondrial dysfunction alters collagen and extracellular matrix gene expression resulting in the connective tissue disorder. Mechanistically, it has been suggested that mitochondria could regulate collagen metabolism via the redox state (20), which is significantly altered in the patient cybrids.

With respect to the underlying cause of the neurological phenotypes in this family, the HDC variant may well be the cause of the Tourette phenotype (7), but the mtDNA variant causes the connective tissue and metabolic phenotypes. This perspective is supported by the fact that the father but not the mother displays a Tourette phenotype and by additional studies in recent years that linked the HDC variant to Tourette syndrome (21, 22). However, the extended neurological manifestations and the skewed inheritance of the HDC gene in the surviving offspring cannot be explained solely by inheritance of the HDC gene. Rather, the mtDNA defects must also contribute to the diverse neurological symptoms of the offspring as well as the connective tissue and metabolic manifestations of the mother and her children. It has already been shown that a mtDNA mutation is sufficient to cause ASD-like symptoms in mice (6) and mitochondrial dysfunction is thought to be a key feature in neurodegenerative disorders (23). RNASeq analysis of the patient transmitochondrial cybrids identified the dysregulation of genes associated with neurodevelopment and axon guidance. In addition, we showed that histamine aggravates the mitochondrial phenotype *in vitro*. Given the prominent role of histaminergic signaling in the brain, it is possible that both the HDC variant as well as the mtDNA variant account for the inheritance of the neuropsychological disorders of the children. The beneficial effects of N-acetylcysteine and nicotinamide riboside on the mitochondrial dysfunction of these patients suggests potential therapeutic options, some of which might be generalizable to other mitochondrially-related neuropsychiatric and/or connective tissue disorders.

## Online Methods

### mtDNA Sequencing

The entire mtDNA was PCR amplified in two fragments using the SequalPrep Long PCR Kit (Invitrogen #A10498) with the following primers: (1) hmtL9611: 5’-TCCCACTCCTAAACACATCC-3’, hmtH1405: 5’-ATCCACCTTCGACCCTTAAG-3’; and (2) hmtL1305: 5’-GTAAGCGCAAGTACCCACG-3’, hmtH9819: 5’-GCCAATAATGACGTGAAGTCC-3’. The PCR products were visualized on an agarose gel with ethidium bromide to assess amplification and product size specificity. For library preparation the two PCR fragments were combined in equal ratios to make 200 ng of DNA, fragmented and blunt-end ligated to 3’ P1 and barcoded 5’ A-BC adaptors (Life Technologies) using the NEBNext Fast DNA Fragmentation & Library Prep Set for Ion Torrent (New England Biolabs #E6285S/L). Size selection was done using Agencourt AMPure XP beads (Beckman Coulter). Ready amplified libraries were quantified with the Ion Library Quantitation kit (Life Technologies #4468802). Multiple barcoded samples per run were pooled in equimolar ratios and templated Ion Sphere Particles (ISPs) were prepared and enriched on the One Touch 2 and ES systems using the Ion PGM Template OT2 200 kit (Life Technologies #4480974). The enriched templated ISPs were then loaded on an Ion 318 sequencing chip and sequenced on an Ion Personal Genome Machine (PGM) using Ion PGM Sequencing 200 v2 kit (Life Technologies). Data was analyzed using the Ion Torrent Suite v4.0.1 and NextGENe software v2.3.3. Haplogroups were identified using the plugin MtDNA Variant Caller 4.4.3.3 and the HaploGrep algorithm.

### Cell lines and treatments

Venous blood samples were obtained from the patient family and three controls following informed consent. Leukocytes were isolated from whole blood by Ficoll-Hypaque gradients and the isolated cells were EBV transformed to establish lymphoblast cell lines by Corriell. Lymphoblasts were maintained in RPMI 1640 medium (11879, Gibco) supplemented with 10% heat-inactivated fetal calf serum, 1 mM glucose, and 2 mM uridine. For all lymphoblastoid cell line experimental procedures, 4 million cells were seeded in 15 ml of culture media 48h prior to the experiment.

Transmitochondrial cybrids were created from the most affected daughter (Fig. 1, II5) and the closest-matching control. Lymphoblasts were enucleated with actinomycin D (0.5 µg/mL) for 14 hrs and then fused with 143B(TK-) ρ^0^ osteosarcoma cells using 45% polyethylene glycol/DMSO. Cells were plated and grown under selection for 4 weeks in DMEM containing 10% dialyzed FBS, 50 µg/mL BrdU, and no uridine (24). Clones were picked and grown separately as individual cell lines, resulting in 2 clones of patient cybrids harboring the patient mtDNA and 1 clone of control cybrids harboring the mtDNA of the closest-matching control. Both patient clones were used for the biochemical and bioenergetic comparison to control cybrids, and showed very similar responses. The results were pooled for statistical and graphical representation.

All cybrids were maintained in DMEM medium (11966, Gibco) supplemented with 10% heat-inactivated fetal calf serum, 1 mM glucose and 2 mM uridine. For all experimental procedures, 2.5 million cells were seeded in T75 cell culture flasks in 10 ml culture media 48 h prior to the experiment.

For histamine treatment, a 100 mM histamine stock solution in water was prepared fresh the day of use 100 µM histamine was added to the cells either 24 h prior or immediately prior to the respective measurements.

For pyrilamine treatment, a 100 mM pyrilamine stock solution in PBS was prepared and stored at -20°C. 100 µM pyrilamine was added to the cells 24 h prior to the respective measurements and before histamine addition.

For thapsigargin treatment, a 1 mM thapsigargin stock solution in DMSO was prepared and stored at -20°C. 2 µM thapsigargin was added to the cells immediately prior to the measurements.

For the MCU inhibitor treatment with KB-R7943, a 5 mM KB-R7943 stock solution in DMSO was prepared and stored at -20°C. 10 µM KB-R7943 was added to the cells 5 min prior to the measurements.

For nicotinamide riboside (NR) and N-acetyl-cysteine (NAC) treatments, the culture media was supplemented with 300 µM NR or 1mM NAC and cells were cultured in the continuous presence of both compounds. Representative measurements were started 1 week after initiation of the treatments and continued for 3 weeks. All parameters assessed were checked for trends coinciding with treatment duration, but cellular physiology remained stable.

### Doubling time

The doubling time was determined in conjunction with the respirometry and flow experiments. Cells were harvested and counted for respirometry/flow cytometry. This allowed calculation of the doubling time via the formula:

Doubling time (h) = culture time (h) / log_2_(cell number_end_ / cell number_start_)

### High Resolution respirometry in intact cells

Respiration in intact cells was measured as described (25). Briefly, the Oroboros Oxygraph-2k was first calibrated to air with the respective cell culture medium of the cells to be measured (block temperature: 37°C, stirrer speed = 750 rpm, oxygen sensor gain = 4, data recording interval 2 sec). Subsequently, the cells were harvested and 2 million cells, resuspended in their conditioned medium, were loaded into each chamber of the respirometer. The first oxygen consumption plateau corresponds to Routine respiration, determining respiration under endogenous substrates and energy demand. Addition of 0.5 µl of 5 mM oligomycin blocks complex V, resulting in Leak respiration, which determines proton leak across the inner mitochondrial membrane independently of complex V. Stepwise addition of 1 mM carbonyl cyanide p-trifluoro-methoxyphenyl hydrazone (FCCP) reveals the electron transport system (ETS) capacity, which determines the maximum respiratory capacity under infinite energy demand. Lastly, 5 µM antimycin A was added to determine background respiration, which is subtracted from the other respiratory states.

Respirometry and parallel ROS production using Amplex UltraRed were performed as described previously (26). Briefly, air calibration was performed using the same settings as for intact cells, but in Mir05 respiratory buffer [0.5 mM EGTA, 3 mM MgCl_2_, 60 mM lactobionic acid, 20 mM taurine, 10 mM KH_2_PO_4_, 20 mM HEPES, 110 mM D-sucrose, and fatty acid-free BSA (1 g/l) and 5 mM DTPA]. After air calibration, the chambers were closed and 10 µM Amplex Ultrared (10 mM stock in DMSO), 0.5 U/ml horseradish peroxidase (500 U/ml stock in Mir05), and 5 U/ml superoxide dismutase (5000 U/ml stock in Mir05) were added. Subsequently, ROS calibration was performed with 2 consecutive injection of 0.2 µM hydrogen peroxide (Fluorescence-sensor Green, gain for sensor = 1000, polarization voltage = 300 mV).

Cells were harvested and 2 million cells in 400 µl Mir05 were added to each chamber. Cells were permeabilized by addition of 5 µg/ml digitonin. Addition of 5 mM pyruvate, 2 mM malate, 10 mM glutamate and 5 mM adenosine diphosphate (ADP) resulted in complex I-linked respiration. Further addition of 10 mM succinate revealed OxPhos capacity, the maximal capacity of mitochondria to produce ATP upon substrate saturation. Addition of 1.25 µM oligomycin resulted in Leak respiration and stepwise addition of FCCP resulted in ETS capacity. Addition of 2.5 µM rotenone blocked complex I and revealed complex II-linked respiration. Lastly, 5 µM antimycin A was added to determine background respiration, which is subtracted from the other respiratory states. ROS production at the different respiratory states were first corrected for background flux before addition of cells and calculated as ROS production/respiration normalized to control.

### Flow Cytometry

For all flow cytometry experiments, cells were seeded 48 h prior in T75 cell culture flasks in their respective medium. The day of the experiment, cells were harvested, counted and for each condition 1 million alive cells were stained in Tyrodes buffer (135 mM NaCl, 5 mM KCl, 1.8 mM CaCl_2_, 20 mM HEPES, 5 mM glucose, 1 mM MgCl_2_, pH 7.4) with the respective dye. Dependent on the dye, cells were washed in Tyrodes buffer, divided into 2 technical replicates, and 10,000 cells were measured for each replicate in Tyrodes buffer on an LSR Fortessa (BD). Alive singlets were gated in the FSC/SSC scatter plot and the mean fluorescence of the respective channel was quantified, corrected for the mean fluorescence of an unstained cell sample in that channel. Mitochondrial localization of the mitochondrial dyes following the respective staining procedures was checked prior to flow cytometry experiments using confocal microscopy.

### Mitochondrial mass using Mitotracker

For determination of mitochondrial mass, cells were stained with 100 nM MitoTracker^TM^ Deep Red FM or 100 nM MitoTracker^TM^ Red CMXRos for 30 min at 37°C. The mean fluorescence was quantified in the APC channel (ex: 640 nm, em: 670/14 nm) for MitoTracker^TM^ Deep Red FM or the mCherry channel (ex:561nm, em:610/20 nm) for MitoTracker^TM^ Red CMXRos.

### Mitochondrial Membrane Potential using JC1

For determination of mitochondrial membrane potential, cells were stained with 1 µg/ml JC-1 for 30 min at 37°C. The mean fluorescence was quantified in the FITC channel (ex:488 nm, em:530/30 nm) and the PE channel (ex:561 nm, em:582/15 nm), and the ratio of PE/FITC was calculated as a surrogate for membrane potential. This was verified by measuring cells treated with 5 µM FCCP upon staining, which showed a lower PE/FITC fluorescence ratio.

### Mitochondrial ROS

For determination of mitochondrial ROS, cells were stained with 5 µM MitoSOX^TM^ Red for 15 min at 37°C. The cells were spun down and resuspended in fresh Tyrodes buffer. The mean fluorescence was quantified in the PerCP channel (ex:488 nm, em:695/40 nm).

### Cytosolic ROS

For determination of cytosolic ROS, cells were stained with 1 µM H2DCFDA for 30 min at 37°C. The mean fluorescence was quantified in the FITC channel (ex:488 nm, em:530/30 nm).

### Mitochondrial calcium

For determination of mitochondrial calcium, cells were stained with 10 µM Rhod-2 AM (1 mM stock in DMSO with potassium borohydride) + 0.02% pluronic acid for 30 min at room temperature. The cells were washed once and resuspended in fresh Tyrodes buffer. The mean fluorescence was quantified in the PE channel (ex:561 nm, em:582/15 nm).

### Cytosolic calcium

For determination of cytosolic calcium, cells were stained with 1 µM Fura Red^TM^ AM + 0.02% pluronic acid for 30min at 37°C. The cells were spun down and resuspended in fresh Tyrodes buffer. The mean fluorescence was quantified in the BV650 channel (ex:405 nm, em:655/8 nm) and the PerCP channel (ex:488 nm, em:695/40 nm). The ratio of BV650/PerCP was calculated as a surrogate for membrane potential. In confocal microscopy, cells were excited at 405 nm and 488 nm and emission was detected between 500 nm to 758 nm.

### Apoptosis

For determination of apoptosis, cells were harvested, washed in DPBS and 0.5 million cells were stained in 200 µl annexin-binding buffer (10 mM HEPES, 140 mM NaCl_2_, 2.5 mM CaCl_2_, pH 7.4) supplemented with either 10 µl Annexin V Pacific Blue^TM^ or 10 µl Annexin V FITC^TM^ for 15 min at room temperature. After staining, 800 µl annexin-binding buffer was added and the cells were measured using flow cytometry. The mean fluorescence was quantified in the Pacific Blue channel (ex:405 nm, em:450/50 nm) or the FITC channel (ex:488 nm, em:530/30 nm), respectively. No live-cell gate was applied and percentage of positive cells was quantified and normalized to control.

### Mitochondrial permeability transition pore (mPTP) opening

Mitochondrial permeability transition pore (mPTP) opening was measured using the calein-cobalt method as described (27). Briefly, cells were stained in Kreb’s Ringer buffer (KRB) (135 mM NaCl, 5 mM KCl, 0.4 mM KH_2_PO_4_, 1 mM MgSO_4_, 20 mM HEPES, 5.5 mM glucose, pH 7.4) supplemented with 1 µM calcein, 200 nM MitoTracker^TM^ Red CMXRos, 2 mM CoCl_2_ and 200 µM sulfinpyrazone for 15 min at 37°C. Cells were washed once in modified KRB (KBR+ 1 mM CaCl_2_), split and resuspended in modified KRB with and without 1 µM ionomycin (positive control), and measured using flow cytometry. The mean fluorescence was quantified in the FITC channel (ex:488 nm, em:530/30 nm) for calcein and the mCherry channel (ex:561 nm, em:610/20 nm) for MitoTracker^TM^ Red CMXRos. mPTP opening was calculated as the mean calcein fluorescence of cells treated with ionomycin divided by the mean calcein fluorescence of untreated cells and then normalized to control cells.

### Confocal microscopy

For all confocal microscopy experiments, 50,000 cybrids were seeded in each quadrant of 4-chambered 35 mm dishes with glass bottom (Cellview, 627870) in their standard culture medium 48 h prior to the measurement. On the day of the experiment, medium was exchanged for the respective staining solution and cells were imaged in Tyrodes buffer at 37°C and atmospheric CO_2_ after a 15 min equilibration in the microscope incubation chamber. For lymphoblasts, the 4-chambered 35 mm dishes with glass bottom were precoated with 2.5 µg/cm^2^ Cell-Tak^TM^ (Corning) for 30 min at 37°C, rinsed with PBS twice and 500,000 cells were seeded into each quadrant in Tyrodes buffer immediately prior to the 15 min incubation in the microscope incubation chamber. Imaging was performed using a Zeiss LSM 710 in combination with the Zen 2012 software for image acquisition and analysis.

### Fluorescence lifetime imaging microscopy (FLIM) of NADH autofluorescence

Fluorescence lifetime imaging of NADH was performed on a laser scanning microscope (Zeiss LSM 710) as described (28). Briefly, NADH was excited using two-photon excitation with a pulsed (80 MHz, 100 fs pulse width) titanium-sapphire laser at a power of <5 mW on the sample. Time-correlated single photon counting (TCSPC) with the hybrid detector HPM-100-40 was performed. The detector was coupled to the NDD port of the LSM 710. FLIM images (512 × 512 pixel) were taken at a temporal resolution of 256 time channels within a pulse period of 12.5 ns. Final settings: 60 sec collection time; ≈ 15 µsec pixel dwell time; 135 × 135 µm^2^ scanning area, Plan-Apochromat 63x/1.40 Oil DIC M27 lense, 730 nm excitation wavelength and 460/50 nm bandpass emission filter. Data were recorded using SPCM 9.8 and subsequently analyzed using SPCImage 8.0 assuming a biexponential decay. Final analysis settings: WLS fit method, lifetime components fixed to 400 ps and 2500 ps for free and protein-bound NADH, square binning of 2, peak threshold adapted to background, shift fixed at a pixel of clear NADH signal. The mean lifetime (τ_mean_) was calculated and fitting of the calculated lifetime curve was confirmed by checking the mean χ^2^, which should be below 1.2. For subcellular analysis, nuclei were selected using region of interests (ROIs) with at least five nuclei analyzed per image. The mitochondria-rich regions were selected using the threshold function.

### RNA Sequencing and analysis

For RNA sequencing, cybrids were seeded and cultured as for biochemical measurements, harvested and a cell pellet of 1.5 million cells was flash frozen and sent to Genewiz (Azenta Life Science, South Plainfield, NJ) for RNA sequencing using Poly-A-primers. Eighteen Human Tourett RNA-sequencing Fastq files, consisting of 6 controls and 12 mutant samples, were processed using the STAR alignment (29) tool and subsequently normalized using the RSEM (30) package based upon the hg38 reference genome (31) and the Gencode version 23 gene annotation (32).

Differential gene expression analysis was performed by comparing each gene in the control group vs the treatment group. The voom procedure (33) was used to normalize the RSEM generated expected counts followed by differential expression testing using R package limma (34) to obtain P-values and LogFC. Specifically, a total of 58581 genes were tested for differential expression between the control and treatment samples. Pathway enrichment was performed on control vs treatment samples using Gene Set Enrichment Analysis (GSEA) (35, 36) version 4.1.0 using a weighted scoring scheme and Hallmark and C2 CP genesets.

All input data and code can be found on github: https://github.com/komalsrathi/tourette-analysis

### Metabolomics

For metabolomics, 1.5 million cybrids were seeded in 10 cm cell culture dishes and cultured using standard culture conditions for 48h. The day of the experiment, cells were transferred on ice into a cold room, medium was removed, cells were rinsed once using ice-cold PBS and liquid nitrogen was poured directly on the cells. The cell slush was scrapped of and transferred into a 1.5 ml tube, that was flash frozen in liquid nitrogen and stored at -80°C until metabolite preparation. A separate aliquot was taken for determination of protein concentration using BCA.

For untargeted LC/MS metabolomics, 50 µL of each thawed cell pellet on ice was aliquoted for untargeted metabolomics, and a 5 µL aliquot of each sample was pooled together and divided to make six 50 µL QC samples. Methanol (200 µL) was added to each sample, vortexed vigorously for 10 sec, and centrifuged at 14000 rpm for 10 min at 4 °C. Then, 2 x 100 µL aliquots (one for reversed phase C18 LC/MS and one for HILIC/MS) of the supernatant were placed into one of two 96 well plates and dried under nitrogen at 30°C. Dried samples were reconstituted in 200 µL of HPLC mobile phases before LC/MS analysis on a Thermo Vanquish UHPLC/Orbitrap ID-X mass spectrometer. Compound Discoverer (Thermo Fisher Scientific) was used to generate PCA plots from the metabolite signals extracted from the raw data files, fold changes, p-values, heat maps, whisker plots, and perform a database search for metabolite identification. Pathway analysis was performed in MetaboAnalyst 5.0 (37), using all metabolites that are significantly (p<0.05) up or down in mutant versus control cybrids.

For targeted LC/MS metabolomics of organic acids and malonyl and acetyl CoA, aliquots (100 µL) of frozen cell lysates were homogenized in equal volumes of acetonitrile/0.6 % formic acid. Metabolites were extracted in cold aqueous/organic solvent mixtures according to validated, optimized protocols in our previously published studies (38, 39). These protocols use cold conditions and solvents to arrest cellular metabolism and maximize the stability and recovery of metabolites. Each class of metabolites was separated with a unique HPLC method to optimize their chromatographic resolution and sensitivity. Quantitation of metabolites in each assay module was achieved using multiple reaction monitoring of calibration solutions and study samples on an Agilent 1290 Infinity UHPLC/6495 triple quadrupole mass spectrometer (39). Raw data were processed using Mass Hunter quantitative analysis software (Agilent). Calibration curves (R2 = 0.99 or greater) were either fitted with a linear or a quadratic curve with a 1/X or 1/X2 weighting.

### Posttranslational modifications of histones

Histone isolation from cybrids was performed as described previously (40). Briefly, cells were lysed in nuclear isolation buffer (NIB, 15 mM Tris, 60 mM KCl, 15 mM NaCl, 5 mM MgCl_2_, 1 mM CaCl_2_, 250 mM sucrose, 1mM DTT, 500 µM AEBSF, 5 nM microcystin, 10 mM sodium butyrate) supplemented with 0.3% NP-40 for 5 min, nuclei were pelleted and washed three times in NIB. Histones were extracted in 0.2 M sulfuric acid for 4 h and precipitated from the sulfuric acid using 20% TCA. Histone precipitate was rinsed using ice-cold acetone+0.1% HCL, air-dried, resuspended in water and the concentration determined using BCA.

For histone mass spectrometry, primary amines and monomethyllysine residues of histones were derivatized using propionic anhydride in acetonitrile and ammonium hydroxide. Histones were dried, digested using trypsin, and then propionylation was performed again to derivatize the newly generated N-termini. Samples were dried then desalted using in-house stagetips. Peptides were separated on an EASY-nLC 1000 using 0.1 % formic acid in water as buffer A, 0.1% formic acid in acetonitrile as buffer B, and C18 as trap and analytical stationary phases (41). Data were acquired on a Thermo Q Exactive using DIA and processed in EpiProfile 2.1 (42).

## Acknowledgements

We would like to thank Shiping Zhang for programming the mtDNA allele co-occurrence tool, enabling the analysis in Fig.S1c. We would further like to express our gratitude to Christopher Petucci at the Metabolomics Core at the University of Pennsylvania for performing and analyzing the global and targeted metabolomics and to the Flow Cytometry Core at CHOP for their training in flow cytometry. Lastly, we would like to thank Niagen® for providing the NR.

## Author Contributions

PMS, DCW designed the study; PMS, LSA, ML, JH, KJ, AB, TY, RM performed the experiments; PMS, KR, KJ, MJ, BAG analyzed the data; KK, CAF, DCW performed the clinical evaluation; PMS and DCW wrote the manuscript. All authors read and approved the manuscript.

---

**Fig. S1.**
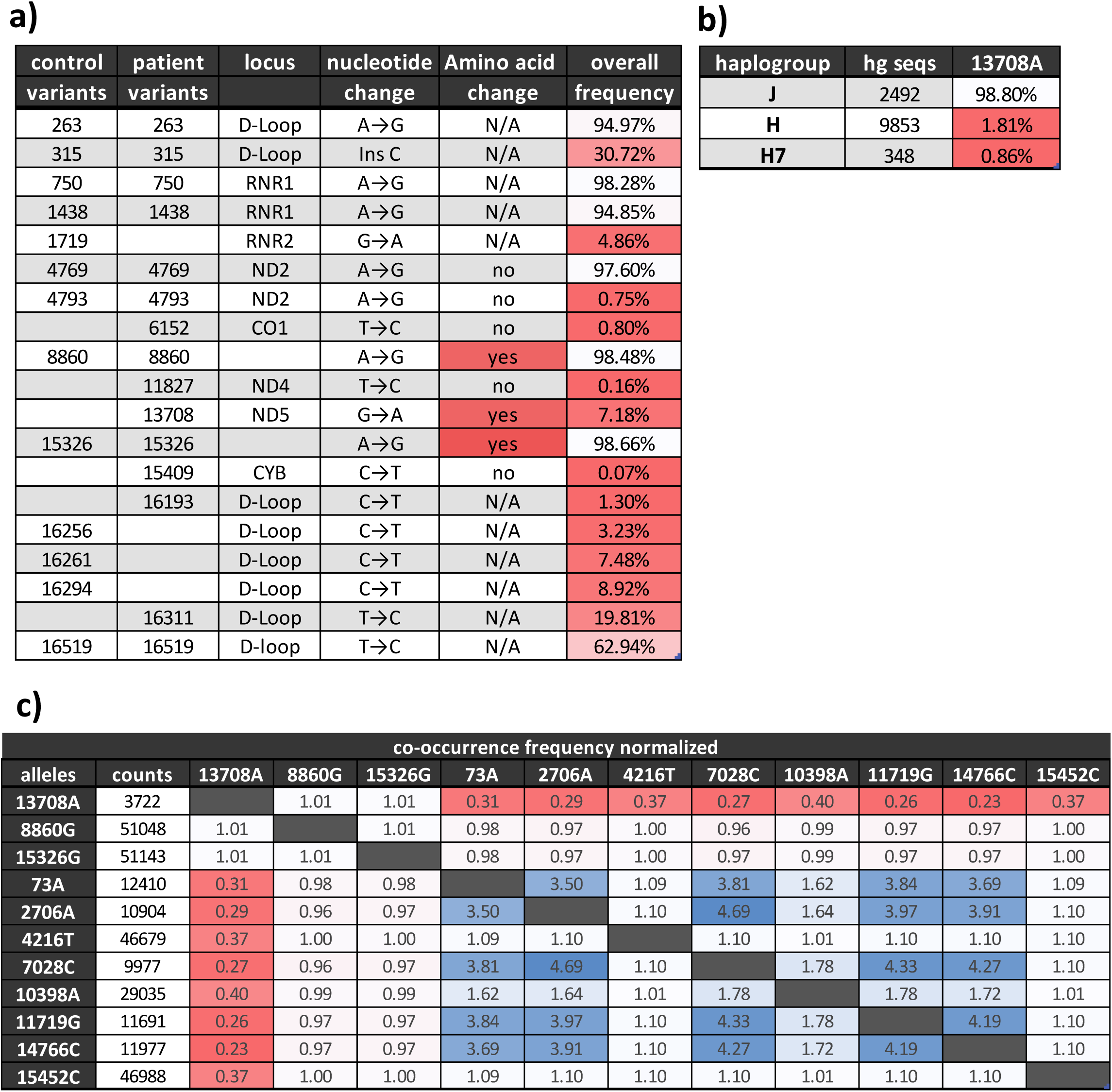
Mitochondrial haplogroup J defining variant *ND5* m.13708G>A on haplogroup H7 background. **a)** Mitochondrial variants found in the proband (mother, I2) and a closely matched control, indicating in which region/gene of the mtDNA the variant is localized, the nucleotide change, the associated amino acid change, and the frequency of the variant on Mitomap (as of 05/04/2021). Variants that result in an amino-acid change are marked in red, frequencies of the variants are colored white to red with red indicating rarer variants. **b)** Frequencies of 13708A in different mtDNA haplogroups colored white to red with red indicating a lower frequency. Counts display the number of mtDNAs of the respective haplogroup on Mitomap (as of 05/04/2021). **c)** Normalized co-occurrence of mtDNA variants with values < 1 (red) indicating lower-than-expected co-occurrence within the population and values >1 (blue) a higher-than-expected co-occurrence (Mitomap, as of 05/04/2021). Co-occurrence is displayed for all proband mtDNA variants and variants between mitochondrial haplogroup H and J that result in an amino acid change.

**Fig. S2.**
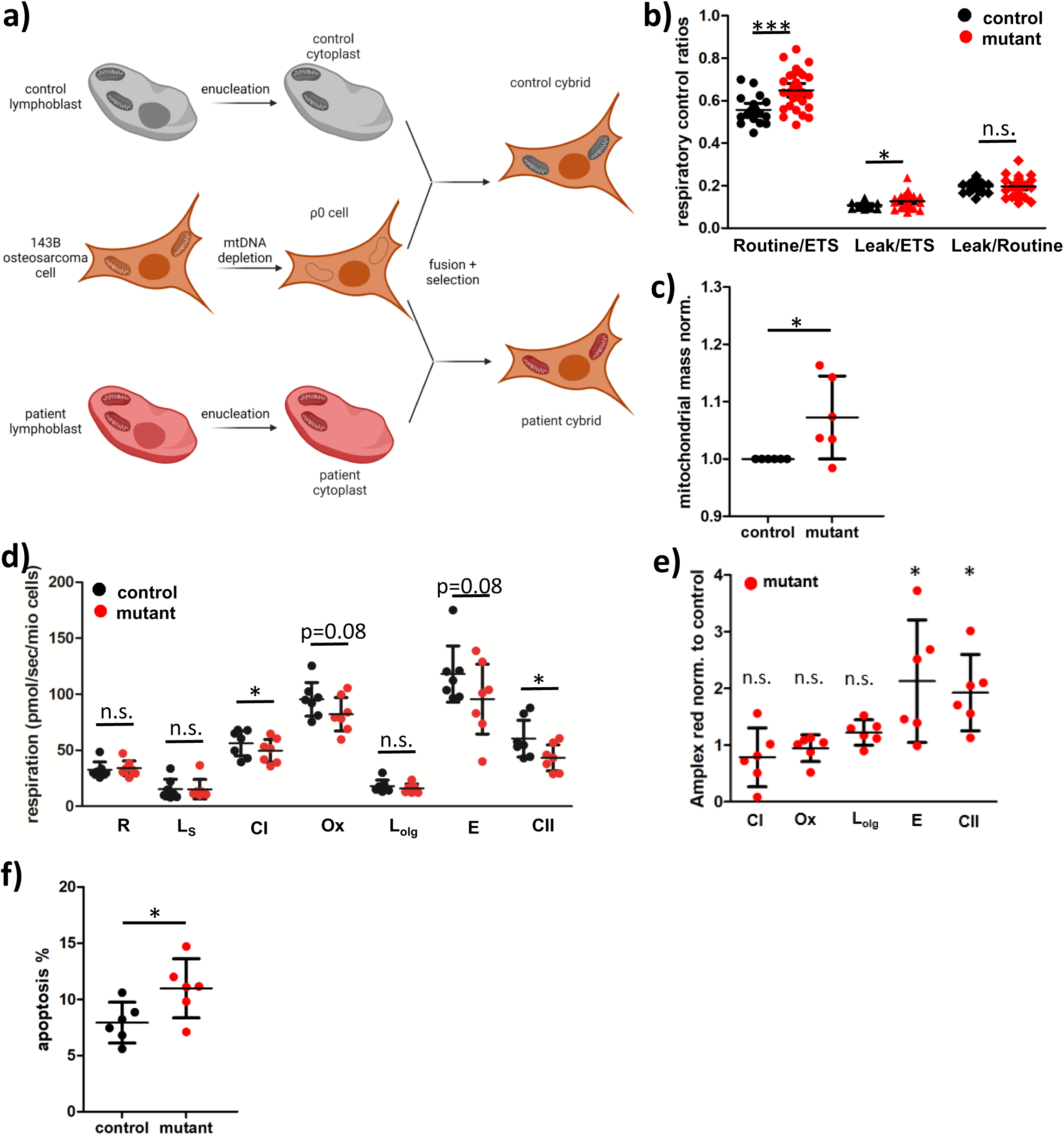
*ND5* m.13708G>A-H7 mtDNA reduces bioenergetic function. **a)** Diagram displaying creation of 143B(TK^-^) transmitochondrial cybrids. Lymphoblasts from control (grey) and proband (Fig. 1a, I2) (red) were enucleated and the resulting cytoplasts fused with ƿ^0^ 143B(TK^-^) cells that are depleted of their mtDNA. After selection, the resulting cybrids contain the 143B(TK^-^) nuclear background with either the patient (mutant) or control mitochondria, allowing the deciphering of the influence of the mtDNA independent of nuclear genes. **b)** Respiratory control ratios of high-resolution respirometry in intact cybrids calculated from data in Fig.1b (Routine/ETS: fraction of maximal respiratory capacity used at baseline; Leak/ETS: fraction of maximal respiratory capacity that is not used for ATP production; Leak/Routine: fraction of baseline respiration that is not used for ATP production) (control: n = 19, mutant n= 31, each in technical duplicates, Mann Whitney test). **c)** Mitochondrial mass of control and mutant cybrids quantified as fluorescence intensity of Mitotracker CMX ROS in flow cytometry normalized to control (n = 6, technical triplicates, paired t-test). **d)** High resolution respirometry in permeabilized cybrids (n = 7, technical duplicates, Wilcoxon signed rank test) **e)** ROS production in permeabilized cybrids quantified as the increase in amplex red fluorescence at different respiratory states normalized to control (n = 6, technical duplicates, One-sample t-test). **f)** Apoptosis in cybrids quantified as the fluorescence intensity of the cells after Annexin V staining using flow cytometry (n = 6, technical duplicates, paired t-test).

**Fig. S3.**
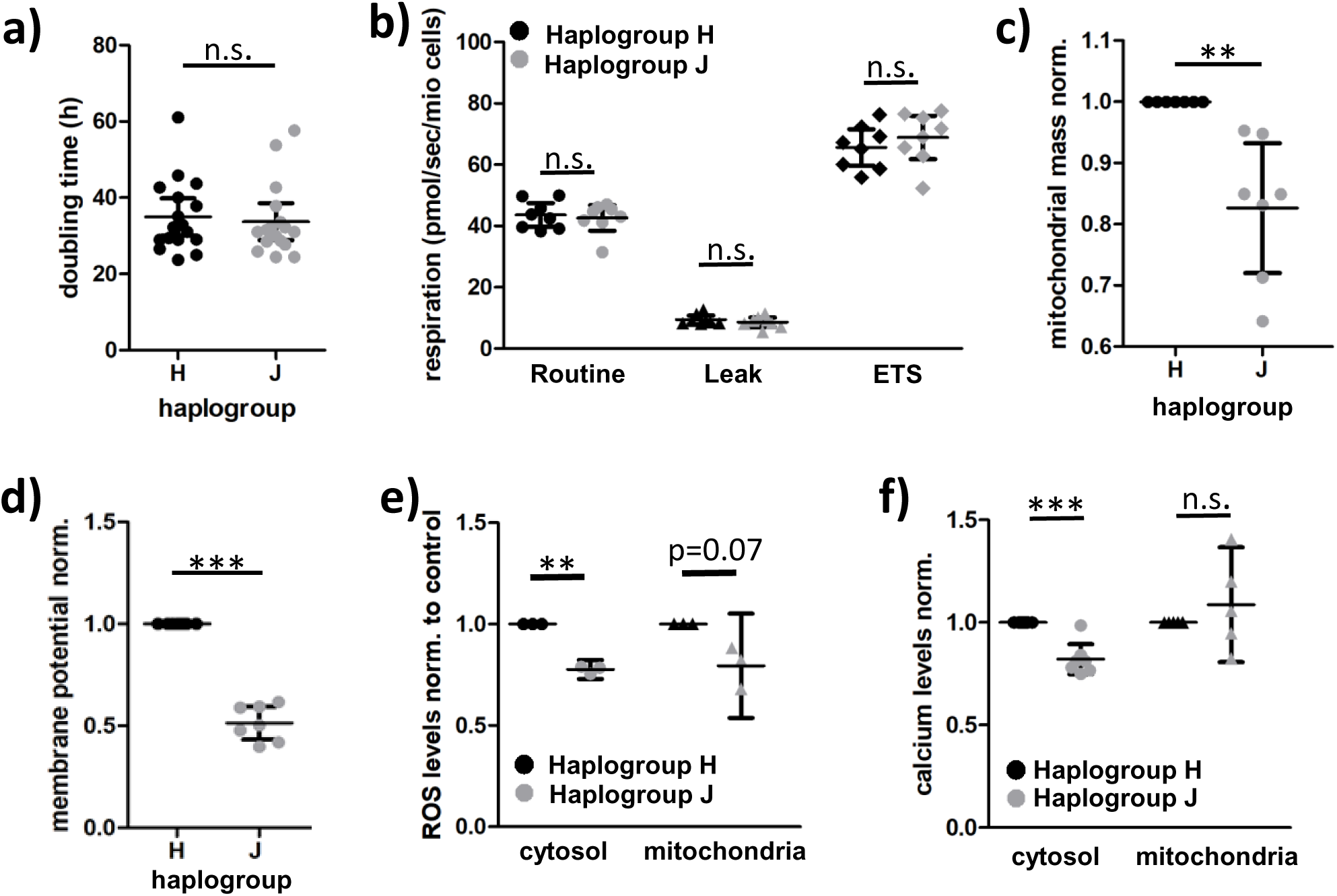
Mitochondrial haplogroup J shows similar bioenergetics to haplogroup H. **a)** Doubling time of haplogroup H and J cybrids in low glucose media (n = 17, Mann Whitney test). **b)** High resolution respirometry of intact haplogroup H (black) or J (grey) cybrids (n = 8, each in technical duplicates, Mann Whitney test). **c)** Mitochondrial mass of haplogroup H and J cybrids quantified as fluorescence intensity of Mitotracker Deep Red in flow cytometry normalized to haplogroup H (n = 6, technical duplicates, paired t-test). **d)** Mitochondrial membrane potential of haplogroup H and J cybrids measured as the ratio of red to green fluorescence of JC-1 quantified by flow cytometry normalized to haplogroup H (n = 7, technical duplicates, paired t-test). **e)** Cytosolic and Mitochondrial ROS levels of haplogroup H and J cybrids quantified as the fluorescence intensity of DCDFA (cytosol, n = 3 in technical duplicates, paired t-test) or Mitosox (mitochondrial, n = 3 in technical duplicates, paired t-test) normalized to haplogroup H. **f**) Cytosolic and Mitochondrial calcium levels of haplogroup H and J cybrids quantified as the fluorescence intensity of Fura Red (cytosol, n=7 in technical duplicates, paired t-test) or Rhod-2 (mitochondrial, n=5 in technical duplicates, paired t-test) normalized to haplogroup H.

**Fig. S4.**
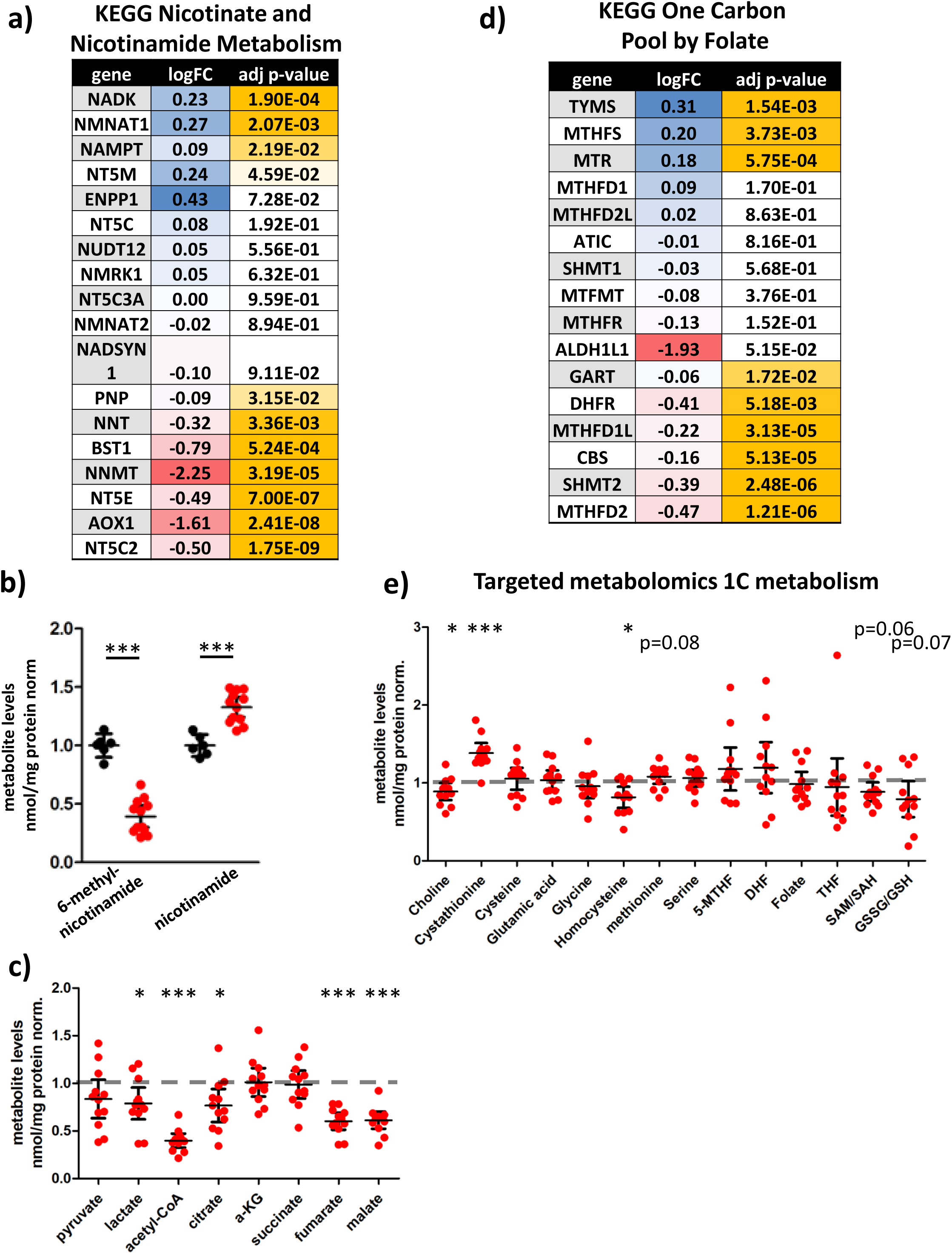
*ND5* m.13708G>A-H7 mtDNA alters nicotinamide, TCA cycle, and one carbon metabolism. **a)** Differential gene expression between mutant and control cybrids for the KEGG Nicotinate and Nicotinamide Metabolism pathway. Log fold change (LogFC) is colored blue – white – red with blue indicating upregulation and red indicating downregulation of a gene in mutant cybrids compared to control. Adjusted (adj.) p-values are colored white – yellow with yellow indicating stronger significance. **b)** Relative nucleotide levels in mutant cybrids normalized to control (n = 6/12 for control/mutant, unpaired t-test) detected in global metabolomics. **c)** Targeted metabolomics of TCA cycle intermediates in mutant cybrids normalized to control (n = 6 for control, n = 12 for mutant, One-sample t-test). **d)** Differential gene expression between mutant and control cybrids for the KEGG One Carbon Pool by Folate pathway. Log fold change (LogFC) is colored blue – white – red with blue indicating upregulation and red indicating downregulation of a gene in mutant cybrids compared to control. Adjusted (adj.) p-values are colored white – yellow with yellow indicating stronger significance. **e)** Targeted metabolomics of one carbon metabolism in mutant cybrids normalized to respective control (n = 6 for control, n = 12 for mutant, One-sample t-test).

**Fig. S5.**
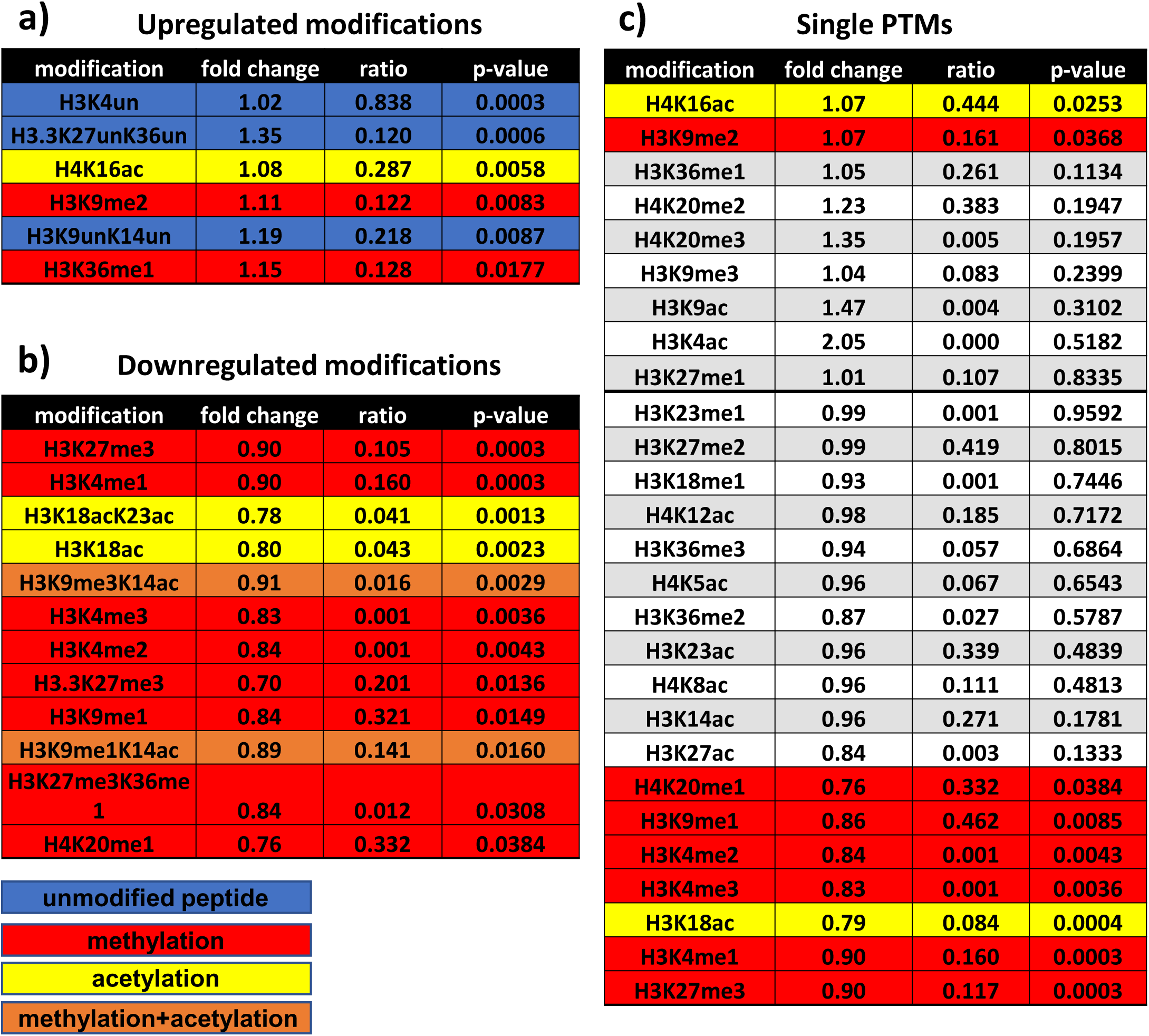
*ND5* m.13708G>A-H7 mtDNA results in histone hypomethylation. **a/b)** Upregulated (a) and downregulated (b) post-translational modifications (PTMs) from histones H3 and H4 in mutant compared to control cybrids measured using mass spectrometry. The ratio indicates the fraction of the peptide with the respective modification, and the color-coding the type of modification present on the peptide, with blue being unmodified, red being methylation, yellow acetylation and orange methylation and acetylation. **c)** Single peptide modifications independent of the originating histone in mutant compared to control cybrids. Significant PTMs are colored according to their modification. Significance versus control was calculated for all PTMs using unpaired t-test (n=6).

**Fig. S6.**
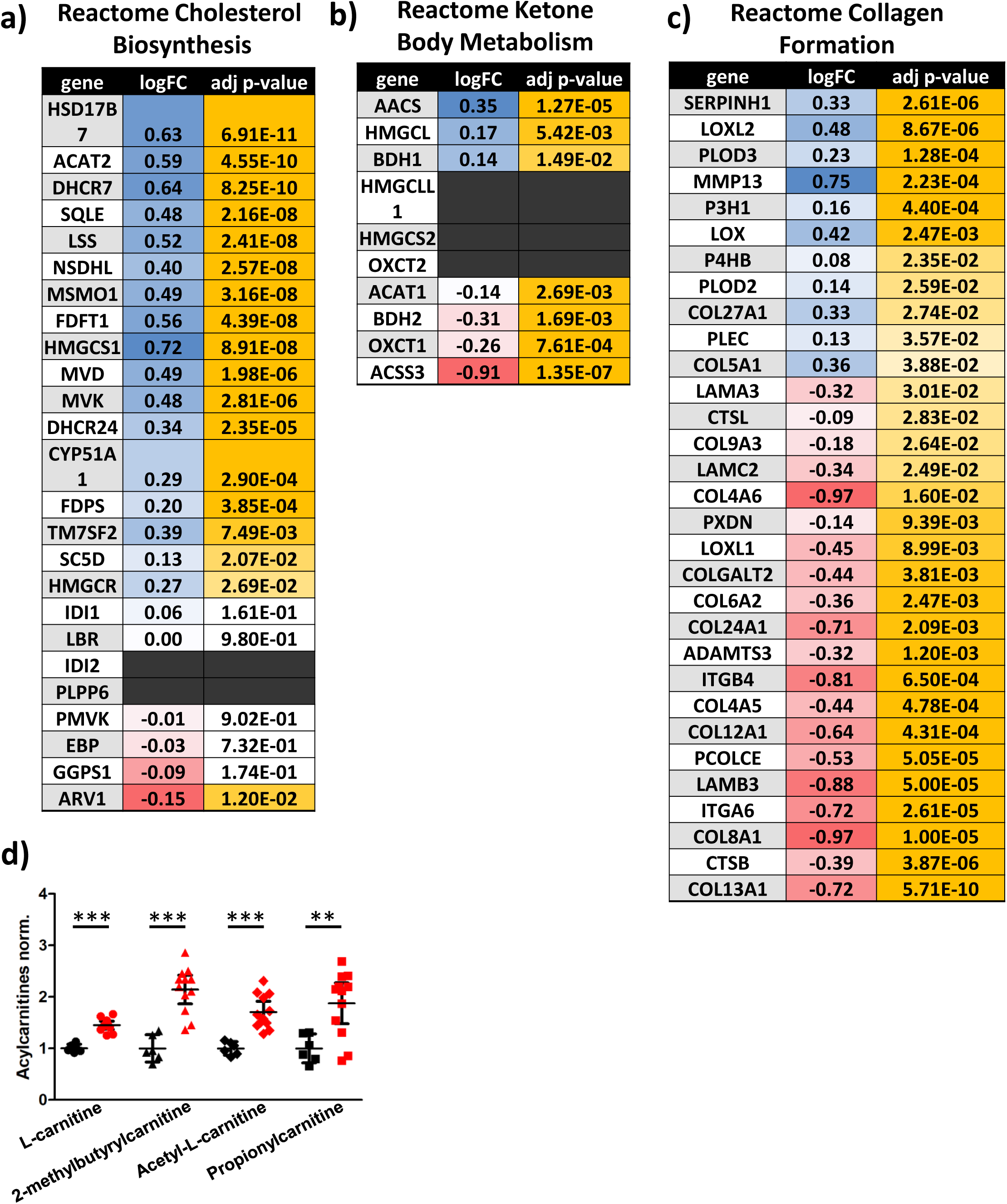
*ND5* m.13708G>A-H7 mtDNA alters cholesterol, ketone and collagen metabolism. **a/b/c)** Differential gene expression between mutant and control cybrids for the Reactome cholesterol biosynthesis (a), the Reactome Ketone Body Metabolism (b) pathway and the Reactome collagen formation pathway (c). Log fold change (LogFC) is colored blue – white – red with blue indicating upregulation and red indicating downregulation of a gene in mutant cybrids compared to control. Adjusted (adj.) p-values are colored white – yellow with yellow indicating stronger significance. **d)** Acylcarnitines detected in the global metabolomics screen in mutant cybrids normalized to control (n = 6 for control, n= 12 for mutant, unpaired t-test).

**Fig. S7.**
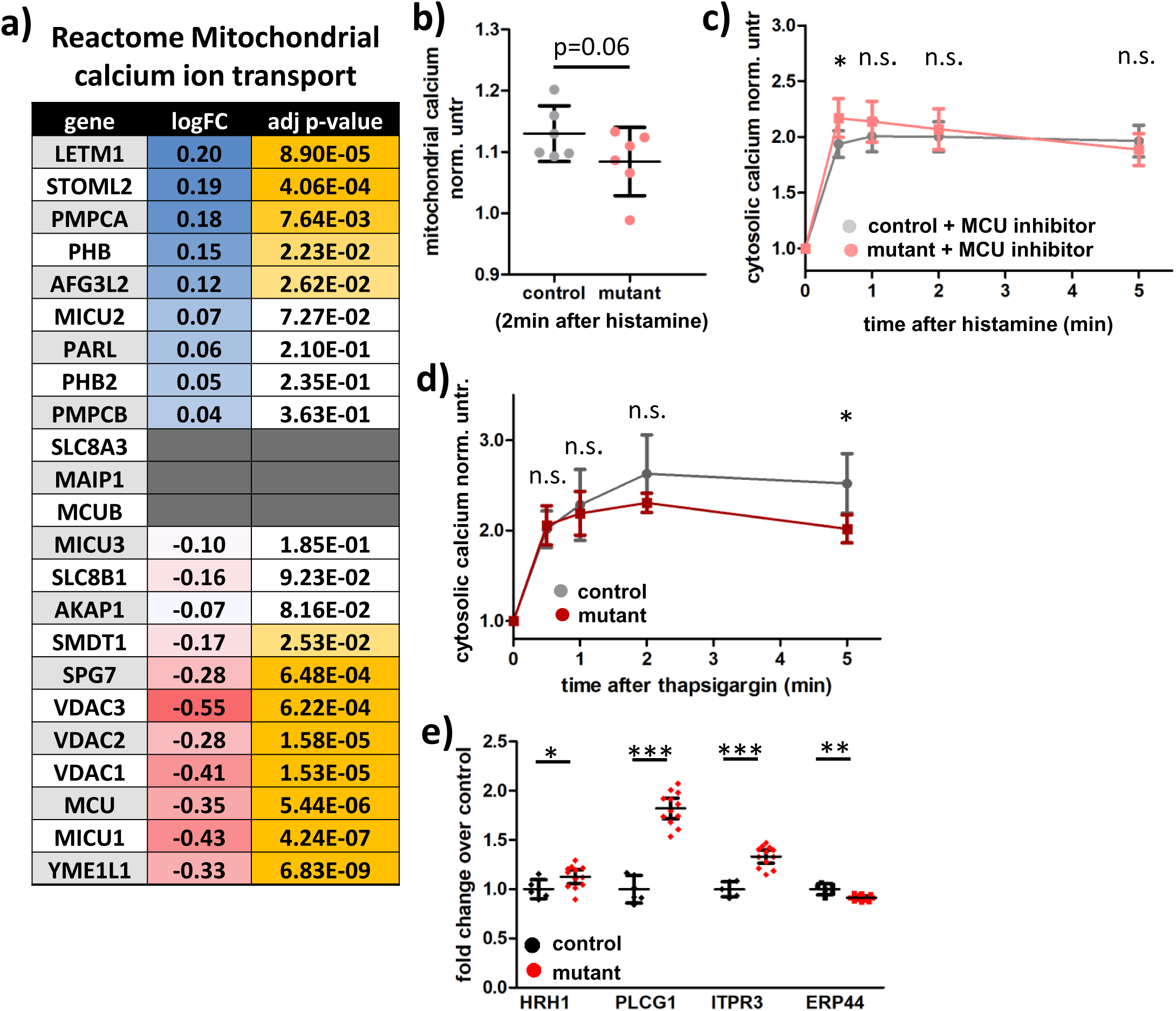
*ND5* m.13708G>A-H7 mtDNA reduces mitochondrial calcium uptake and increases histamine sensitivity. **a)** Differential gene expression between mutant and control cybrids for the Reactome mitochondrial calcium ion transport pathway. Log fold change (LogFC) is colored blue – white – red with blue indicating upregulation and red indicating downregulation of a gene in mutant cybrids compared to control. Adjusted (adj.) p-values are colored white – yellow with yellow indicating stronger significance. **b)** Mitochondrial calcium levels in mutant and control cybrids 2 min after a 100 µM Histamine stimulus, normalized to calcium levels pre-histamine. Measured using Rhod-2 and flow cytometry (n = 6, each in technical duplicates, paired t-test). **c)** Cytosolic calcium levels in variant and control cybrids in response to a 100 µM histamine stimulus after pretreatment with MCU-inhibitor (10 µM KB-R7943), normalized to calcium levels pre-histamine (time point 0). Measured using Fura Red and confocal microscopy (n = 9, paired t-test). **d)** Cytosolic calcium levels in mutant and control cybrids in response to a 2 µM thapsigargin stimulus, normalized to calcium levels pre-thapsigargin (time point 0). Measured using Fura Red and confocal microscopy (n = 6, paired t-test). e) Differential gene expression between mutant and control cybrids for histamine 1 receptor (HRH1), phospholipase C (PLCG1) and IP3 receptor (ITPR3) and endoplasmatic reticulum protein 44 (ERP44), a negative regulator of IP3 receptor-mediated calcium release (n = 6 for control, n = 12 for mutant, unpaired t-test).

**Fig. S8.**
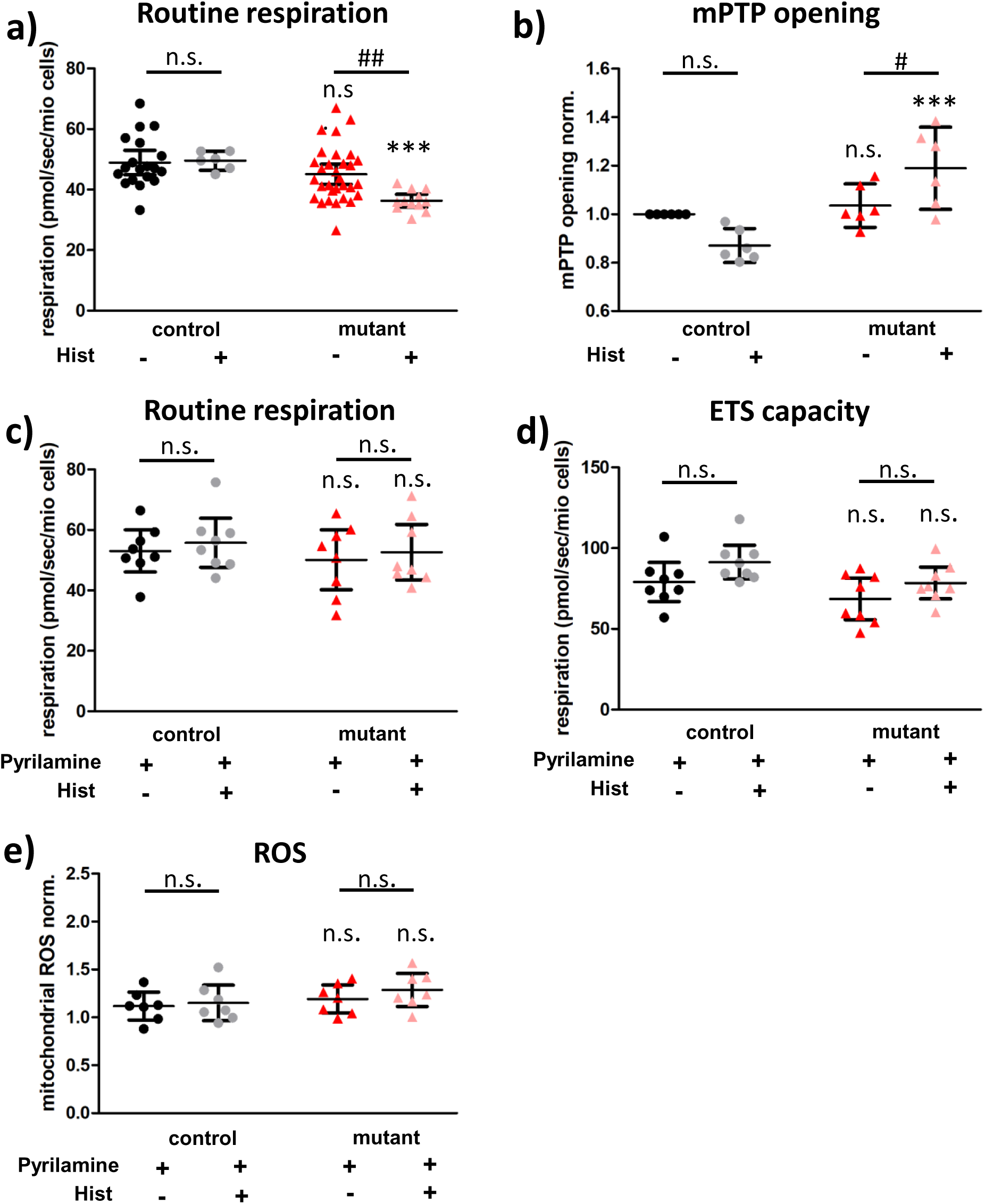
Histamine toxicity in *ND5* m.13708G>A-H7 mtDNA cybrids is mediated via Histamine 1 receptor-mediated calcium release and mPTP opening. **a)** Routine respiration of intact control and mutant cybrids with and without pretreatment with 100 µM histamine for 24h measured using high-resolution respirometry (n = 19, 6, 31, 12 from left to right, each in technical duplicates, Kruskal Wallis test). **b)** mPTP opening in control and mutant cybrids with and without pretreatment with 100 µM histamine for 24h measured cobalt-calcein quenching in flow cytometry (n = 6, each in technical duplicates, One-Way ANOVA). **c/d)** Routine respiration (c) and electron transport system capacity (d) of intact pyrilamine-treated (100 µM) control and mutant cybrids with and without treatment with 100 µM histamine for 24 h measured using high-resolution respirometry (n=8, each in technical duplicates, Kruskal Wallis test). **e)** Mitochondrial ROS levels of pyrilamine-treated control and variant cybrids with and without treatment with 100 µM histamine for 24h quantified as the fluorescence intensity of Mitosox normalized to untreated control (n = 7, each in technical duplicates, One-way ANOVA). Significances between mutant and control are indicated by * and between treatment and control are indicated by #.

**Fig. S9.**
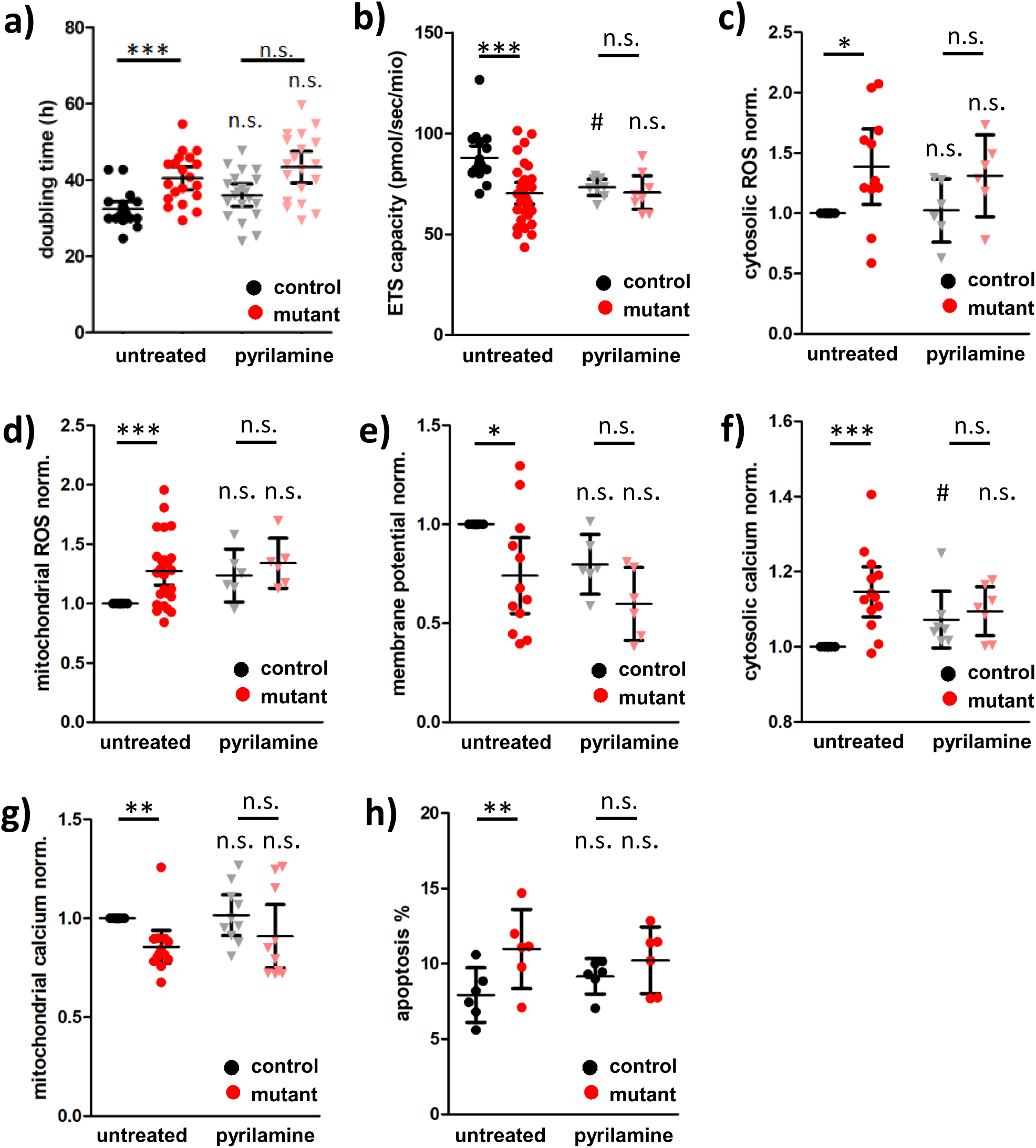
Histamine 1 receptor inhibition reduced mitochondrial function but alleviates the negative effect of the *ND5* m.13708G>A-H7 mtDNA. Comparison of control and mutant cybrids untreated or cultured with 100 µM pyrilamine added to the medium. **a)** Doubling time (n=20, Kruskal Wallis test). **b)** Electron transport system capacity of intact cells (n = 19, 31, 8, 8 from left to right, each in technical duplicates, One-Way ANOVA). **c/d)** Cytosolic (c) and mitochondrial (d) ROS levels quantified as the fluorescence intensity of DCDFA (cytosol, n = 11, 11, 6, 6 from left to right, in technical duplicates, One-way ANOVA) or Mitosox (mitochondrial, n = 26, 26, 6, 6 from left to right, in technical duplicates, One-way ANOVA) normalized to untreated control. **e)** Mitochondrial membrane potential measured as the ratio of red to green fluorescence of JC-1 quantified by flow cytometry normalized to untreated control (n = 12, 12, 6, 6 from left to right, each in technical duplicates, One-way ANOVA). **f/g)** Cytosolic (f) and mitochondrial (g) calcium levels measured by flow cytometry using Fura Red (cytosol, n = 13, 13, 7, 7 from left to right, each in technical duplicates, Kruskal-Wallis test) or Rhod-2 (mitochondria, n = 13, 13, 10, 10, each in technical duplicates, Kruskal-Wallis test) normalized to untreated control. **h)** Apoptosis quantified as the fluorescence intensity of the cells after Annexin V staining using flow cytometry (n = 6, technical duplicates, Repeated measures ANOVA). Significances between mutant and control are indicated by * and between treatment and control are indicated by #.

**Fig. S10.**
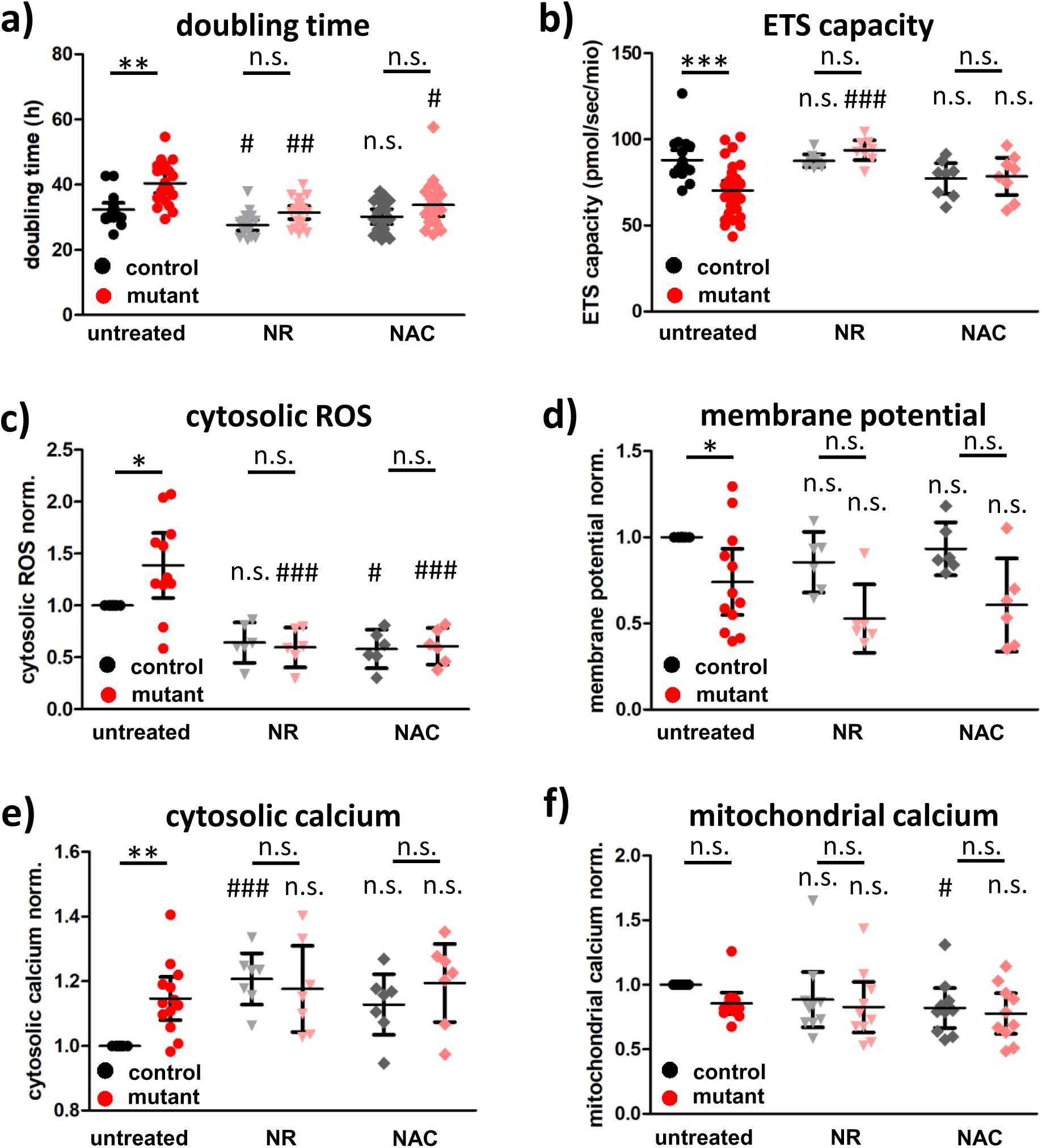
NR and NAC can partially rescue the mitochondrial defect in the *ND5* m.13708G>A-H7 mtDNA cybrids. Comparison of control and mutant cybrids untreated or cultured with either 300 µM nicotinamide riboside or 1 mM N-acetylcysteine added to the medium. **a)** Doubling time (n = 20, Kruskal Wallis test). **b)** Electron transport system capacity of intact cells (n = 19, 31, 8, 8, 8, 8 from left to right, each in technical duplicates, One-Way ANOVA). **c)** Cytosolic ROS levels quantified as the fluorescence intensity of DCDFA (cytosol, n = 11, 11, 6, 6, 6, 6 from left to right, in technical duplicates, One-way ANOVA) normalized to untreated control. **d)** Mitochondrial membrane potential measured as the ratio of red to green fluorescence of JC-1 quantified by flow cytometry normalized to untreated control (n = 12, 12, 6, 6, 6, 6 from left to right, each in technical duplicates, One-way ANOVA). **e/f)** Cytosolic (e) and mitochondrial (f) calcium levels measured by flow cytometry using Fura Red (cytosol, n = 13, 13, 7, 7 from left to right, each in technical duplicates, Kruskal-Wallis test) or Rhod-2 (mitochondria, n = 13, 13, 10, 10, each in technical duplicates, Kruskal-Wallis test) normalized to untreated control. Significances between mutant and control are indicated by * and between treatment and control are indicated by #.

## Detailed clinical findings

Summation of previous report (1) with augmented familial observations.

**I1: Male, age 48**: (Tourette syndrome, Obsessive-Compulsive Disorder [OCD]). “history of motor tics including shoulder shrugs and neck stretching. Vocal tics included persistent clearing of the throat” (1).

**I2: Female, age 46** (connective tissue, metabolic alterations). “She has no history of tics, OCD, trichotillomania, or Attention Deficit Disorder (ADHD)” (1). Additional familial reported findings: pseudotumor cerebri (increased pressure around brain), joint hypermobility with post-compression detethering and decompression (tethered cord syndrome: tissue attachments that limit the movement of the spinal cord within the spinal column), chiari malformation (extension of brain into spinal cord, cranio-cervical mobility requiring fusion), hypermobility below cranio-cervical fusion, chronic diarrhea, ketosis, difficulties with temperature regulation, weight loss and nausea on Diamox (carbonic anhydrase inhibitor).

**II1: Female, age 25** (Tourette syndrome, OCD, metabolic alterations). Aspergers (autism spectrum disorder), ADHD, OCD, hair pulling (trichotillomania), skin-picking (dermatillomania) (1). Additional familial reported findings: Low energy and frequent mild fevers.

**II2: Male, age 23** (Tourette syndrome, connective tissue, and metabolic alterations). “Motor tics…” “Vocal tics consisted of throat clearing”. “Type I diabetes, severe myopia, symptomatic chiari malformation, and tethered spinal cord”, “joint hypermobility,..high, narrow palate..” “tricuspid and mitral valve” dysfunction, “diagnosed with a variant of EDS [Ehler-Danlos syndrome]” (1). Additional familial reported findings: skin herniation which spontaneously improved, possible ankylosing spondylitis (altered posture), kidney issues, DMI (diabetes mellitus insipidus).

**II3: Male, age 21** (Tourette syndrome, connective tissue, and cardiovascular disorders). “mild clubbed feet”, mild joint hypermobility”, ‘chiari malformation”, “myopia, bicuspid aortic valve and root dilation”, “sleep apnea” (1). Additional familial reported findings: Severe cranio-cervical instability, brainstem surgery, aortic aneurism (44mm) with oval aorta on MRI, syncope (loss consciousness) with exercise, supraventricular tachycardia and bradycardia, neurogenic bladder, and patches on retina.

**II4: Male, age 18** (Tourette syndrome, connective tissue, and metabolic alterations). “cardiac involvement, pectus excavatum and high arced palate” (1). Additional familial reported findings: chiari malformation, dysautonomia, and tall (6’3’’).

**II5: Female, age 16** (Tourette syndrome, connective tissue, metabolic and immunological alterations). “Type I diabetes, autonomic dysfunction (tachycardia and narrowed vision), severe joint pain, and symptomatic chiari malformation”, “diagnosed with early-onset osteo-arthritis and a variant of EDS” (1). Additional familial reported findings: post-compression detethering and decompression, muscle weakness, kidney reflux, cranio-cervical instability, ankylosing spondylitis requiring Lidocaine infusions into spinal cord (assessment “life stinks”), dysautonomia, dehiscence (wound reopening post-surgery), unusual blood vessels brain to pelvis with stroke at age five, altered prothrombin time, multiple drug allergies, multiple infections including methicillin-resistant Staphylococcus aureus, Herpes, and Candida.

**II6: Male, age 13** (Tourette syndrome, connective tissue, and metabolic alterations). “evaluated for probable connective tissue disorder with pectus excavatum and hypermobile joints.”: Brain MRI revealed chiari malformation and hydromyelia syrinx”, “surgical … spinal cord detethering”, “ketone utilization disorder”(1). Additional familial reported findings: episodic bradycardia, 3-β-keto-thiolase deficiency, high prolactin levels, carnitine bicitrate therapy.

**II7: Male, age 10**, monozygotic twin (Tourette syndrome and OCD, connective tissue, and metabolic alterations). “symptomatic chiari malformation”, “high/arched palate, bilateral dislocation of hips, and moderate joint laxity”, “mild convex cervical scoliosis centered at C6/C7”, “sleep apnea and breathing pauses while awake”, “possible connective tissue disorder at 6 years of age”, “an EDS variant” (1). Additional familial reported findings: pseudotumor cerebri (increased pressure around brain) treated with ventriculoperitoneal shunt, left eye ptosis, joint hypermobility, post-compression detethering, and decompression, frequently falls asleep, chronically elevated CO2 while sleeping, apnea-hypoapea greater than 10 second, altered prothrombin time.

**II8: Male, age 10,** monozygotic twin (Tourette syndrome, connective tissue, cardiovascular and metabolic alterations).” described as having thin skin, easy bruisability, and joint laxity and dislocation. He has been diagnoses with a tethered spinal cord but has not required surgery”, “bilateral strabismus with surgical correction” (1). Additional familial reported findings: prediabetic (blood sugar 110-112 mg/dL); hypermobile joints with subluxation (partial bone dislocation on CT) of knees, hips, and ankles with severe foot and leg pain; superficial veins on left arm; altered prothrombin time; left ventricular hypertrophy and hypertension starting at 8 years.

